# Spatial soft sweeps: patterns of adaptation in populations with long-range dispersal

**DOI:** 10.1101/299453

**Authors:** Jayson Paulose, Joachim Hermisson, Oskar Hallatschek

## Abstract

Adaptation in extended populations often occurs through multiple independent mutations responding in parallel to a common selection pressure. As the mutations spread concurrently through the population, they leave behind characteristic patterns of polymorphism near selected loci—so-called soft sweeps—which remain visible after adaptation is complete. These patterns are well-understood in two limits of the spreading dynamics of beneficial mutations: the panmictic case with complete absence of spatial structure, and spreading via short-ranged or diffusive dispersal events, which tessellates space into distinct compact regions each descended from a unique mutation. However, spreading behaviour in most natural populations is not exclusively panmictic or diffusive, but incorporates both short-range and long-range dispersal events. Here, we characterize the spatial patterns of soft sweeps driven by dispersal events whose jump distances are broadly distributed, using lattice-based simulations and scaling arguments. We find that mutant clones adopt a distinctive structure consisting of compact cores surrounded by fragmented “haloes” which mingle with haloes from other clones. As long-range dispersal becomes more prominent, the progression from diffusive to panmictic behaviour is marked by two transitions separating regimes with differing relative sizes of halo to core. We analyze the implications of the core-halo structure for the statistics of soft sweep detection in small genomic samples from the population, and find opposing effects of long-range dispersal on the expected diversity in global samples compared to local samples from geographic subregions of the range. We also discuss consequences of the standing genetic variation induced by the soft sweep on future adaptation and mixing.

## I. INTRODUCTION

Rare beneficial alleles can rapidly increase their frequency in a population in response to a new selective pressure. When adaptation is limited by the availability of mutations, a single beneficial mutation may sweep through the entire population in the classical scenario of a “hard sweep”. However, populations may exploit a high availability of beneficial mutations due to standing variation, recurrent new mutation, or recurrent migration [1, 2, 3, 4, 5] to respond quickly to new selection pressures. As a result, multiple adaptive alleles may sweep through the population concurrently, leaving genealogical signatures that distinguish them from hard sweeps. Such events are termed *soft sweeps*. Soft sweeps are now known to be frequent and perhaps dominant in many species [6, 7]. Well-studied examples in humans include multiple origins for the sickle cell trait which confers resistance to malaria [8], and of lactose tolerance within and among geographically separated human populations [9, 10].

Soft sweeps rely on a supply of beneficial mutations on distinct genetic backgrounds, which has two main origins. One is when selection acts on an allele which has multiple copies in the population due to standing genetic variation—a likely source of soft sweeps when the potentially beneficial alleles were neutral or only mildly deleterious before the appearance of the selective pressure [3]. In this work, we focus on the other important scenario of soft sweeps due to recurrent new mutations which arise after the onset of the selection pressure. Soft sweeps become likely when the time taken for an established mutation to fix in the entire population is long compared to the expected time for additional new mutations to arise and establish. In a panmictic population, the relative rate of the two processes is set primarily by the rate at which new mutations enter the population as a whole [5].

Most examples of soft sweeps in nature, however, show patterns consistent with arising in a geographically structured rather than a panmictic population [7]. Spatial structure promotes soft sweeps [11]: when lineages spread diffusively (i.e. when offspring travel a restricted distance between local fixation events), a beneficial mutation advances as a constant-speed wave expanding outward from the point of origin, much slower than the logistic growth expected in a well-mixed population. Therefore, fixation is slowed down by the time taken for genetic information to spread through the range, making multi-origin sweeps more likely. However, the detection of such a *spatial soft sweep* crucially depends on the sampling strategy: the wavelike advance of distinct alleles divides up the range into regions within which a single allele is predominant. If genetic samples are only taken from a small region within the species’ range, the sweep may appear hard in the local sample even if it was soft in the global range.

Between the two limits of wavelike spreading and panmictic adaptation lies a broad range of spreading behaviour driven by dispersal events that are neither local nor global. Many organisms spread through long-range jumps drawn from a probability distribution of dispersal distances (dispersal kernel) that does not have a hard cutoff in distance but instead allows large, albeit rare, dispersal events that may span a significant fraction of the population range [12, 13]. A recent compilation of plant dispersal studies showed that such so-called “fat-tailed” kernels provided a good statistical description for a majority of data sets surveyed [14]. Fat-tailed dispersal kernels accelerate the growth of mutant clones, whose sizes grow faster-than-linearly with time and ultimately overtake growth driven by a constant-speed wave [12, 15]. Besides changing the rate at which beneficial alleles take over the population, long-range dispersal also breaks up the wave of advance [16]: the original clone produces geographically separated satellites which strongly influence the spatial structure of regions taken over by distinct alleles.

Despite its prominence in empirically measured dispersal behaviour and its strong effects on mutant clone structure and dynamics, the impact of long-range dispersal on soft sweeps is poorly understood. Past work incorporating fat-tailed dispersal kernels in spatial soft sweeps [11] relied on deterministic approximations of the jump-driven spreading behaviour of a single beneficial allele [12]. However, recent analysis has shown that deterministic approaches are accurate only in the two extreme limits of local (i.e. wavelike) and global (i.e. panmictic) spreading, and break down over the entire regime of intermediate long-range dispersal [17]. Away from the limiting cases, the correct long-time spreading dynamics is obtained only by explicitly including rare stochastic events which drive the population growth. Deterministic approaches also do not account for the disconnected satellite structure, which has consequences for soft sweep detection in local samples.

Here, we study soft sweeps driven by the stochastic spreading of alleles via long-range dispersal. We perform simulations of spatial soft sweeps in which beneficial alleles spread via fat-tailed dispersal kernels which fall off as a power law with distance, focusing on the regime in which multiple alleles arise concurrently. We find that long-range dispersal gives rise to distinctive spatial patterns in the distribution of mutant clones. In particular, when dispersal is sufficiently long-ranged, mutant clones are discontiguous in space, in contrast to the compact clones expected from wavelike spreading models. We identify qualitatively different regimes for spatial soft sweep patterns depending on the tail of the jump distribution. We show that analytical results for the stochastic jump-driven growth of a *solitary* allele [17], combined with a mutation-expansion balance relevant for spatial soft sweeps [11], allow us to predict the range sizes beyond which soft sweeps become likely. We also analyze how stochastic aspects of growth of independent alleles, particularly the establishment of satellites disconnected from the initial expanding clone, influence the statistics of observing soft sweeps in a small sample from the large population. We find that long-range dispersal has contrasting effects on the likelihood of soft sweep detection, depending on whether the population is sampled locally or globally.

## II. RESULTS

### A. Model of spatial soft sweeps

We consider a haploid population that lives in a *d-*dimensional habitat consisting of demes that are arranged on an integer lattice (e.g. square lattice in *d* = 2). Local resource limitation constrains the deme population to a fixed size 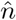, assumed to be the same for all demes. Denoting the linear dimension of the lattice as *L*, the total population size is 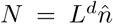. The population is panmictic within each deme. With a rate *m* per generation, individuals migrate from one deme to another. For each dispersal event, the distance *r* to the target deme is chosen from a probability distribution with weight *J*(*r*), appropriately discretized, with the normal-ization 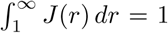. The function *J*(*r*) is called the jump kernel. The dispersal direction is chosen uniformly at random from the unit sphere in *d* dimensions. New mutations arise in all demes at a constant rate *u* per individual per generation. Each new mutation is distinguishable from previous mutations (e.g. due to different genomic backgrounds), but all mutations confer the same selective advantage *s*. Back mutations are ignored. To minimize the effect of the specific boundary geometry, periodic boundary conditions are assumed.

To focus on the effects of long-range dispersal over local dynamics, we now impose a set of bounds on the individual-based parameters following Ref. 11. In particular, we consider only situations where 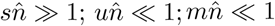 (strong selection, and low mutation and migration rates at the deme level). Mutations are also assumed to be fully redundant, i.e. a second mutation confers no additional advantage. The strong selection condition implies that genetic drift within a deme is irrelevant relative to selection: a new mutation, upon surviving stochastic drift and fixing within a deme (which happens with probability 2*s*) cannot be subsequently lost due to genetic drift. The bounds on mutation and migration rates meanwhile imply that the fixation dynamics of a beneficial mutation within a deme is fast compared to the dynamics of mutation within a deme or of migration among demes. The time to fixation of a beneficial allele from a single mutant individual in the deme, 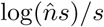, is a few times 1*/s*. When 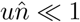 and 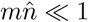, the fixation time scale is much shorter than the establishment time scales of new alleles arising due to mutation or migration, which are 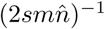 and 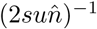 respectively. Therefore, the first beneficial allele that establishes in a deme, whether through mutation or migration, fixes in that deme without interference from other alleles. Furthermore, the assumption of mutual redundancy means that subsequent mutations that arrive after the first fixation event also have no effect. As a result, the first beneficial allele that establishes in a deme excludes any subsequent ones—a situation termed allelic exclusion [11].

Taken together, these assumptions lead to a simplified model that ignores the microscopic dynamics of mutations within demes. For each deme, we keep track of a single quantity: the allelic identity (whether wildtype or one of the unique mutants that has arisen) that has fixed in the deme. At the deme level, new mutations fix within wildtype demes at the rate 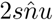, and each mutated deme sends out migrants at rate 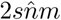 with the target deme selected according to the dispersal kernel *J*(*r*) (the rates explicitly include the fixation probability 2*s* of a single mutant in a wildtype deme). The first successful mutant to arrive at a wildtype deme, whether through mutation or migration, immediately fixes within that deme. The state of the deme thereafter is left unchanged by mutation or migration events, because of allelic excusion.

When time is measured in units of the expected interval 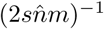 between successive dispersal events per deme, the reduced model is characterized by just three quantities: *L*; *J*(*r*); and the per-deme rate of mutations per dispersal attempt 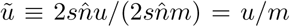, which we call the rescaled mutation rate of our model. Simulations are begun with a lattice of demes of size *L*^*d*^ all occupied by the wildtype. Each discrete simulation step is either a mutation or an attempted migration event, with the relative rates determined by 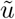 and the fraction of wildtype sites at that step. Mutation events flip a randomly-selected wildtype deme into a new allelic identity. Migration events first pick a mutated origin and then pick a target deme according to the jump kernel. If the target site is wildtype, it acquires the allelic identity of the origin; otherwise the migration is unsuccessful. Simulations are run until all demes have been taken over by mutants.

The fat-tailed jump kernels we use are of the form *J*(*r*) = *μr*^−(1+*μ*)^, with *μ* > 0 to ensure that the kernel is normalizable. The exponent *μ* characterizes the “heaviness” of the tail of the distribution. We have chosen power-law kernels because they span a dramatic range of outcomes that connect the limiting cases of well-mixed and wavelike growth upon varying a single parameter. The growth dynamics of more general fat-tailed kernels in the stochastic regime of interest (i.e. driven by rare long jumps) are largely determined by the power-law falloff of the tail, and details of the dispersal kernel at shorter length scales are less consequential. Therefore, our qualitative results should extend to kernels sharing the same power law behaviour of the tail, provided the typical clones are large enough so that rare jumps picked from the tail of the distribution become relevant. The underlying analysis leading to the results is even more general, and can be applied to any jump kernel that leads to faster-than-linear growth in the extent of an individual clone with time.

The output of a simulation at a given set of *L*, *μ* and 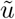 values is the final configuration of mutants, which can be grouped into distinct clones of the same allelic identity. Note that we have ignored the post-sweep mixing of alleles which are now relatively neutral to each other due to migration; this is justified by the separation of time scales between fast fixation and slow neutral migration [11]. In addition, although we restrict ourselves to weak mutation and migration at the deme level, the population-level mutation and migration rates *Nu*, *Nm* are typically large which allows for soft sweeps with strong migration effects.

While our theoretical results are valid for all dimensions, computational limitations prevented us from running extensive simulations in dimensions higher than one. Therefore, we primarily report simulations of linear habitats (*d* = 1) in the main text. Preliminary results from planar simulations (*d* = 2) are reported in Appendix B and are consistent with our theoretical arguments, although quantitative comparisons are limited by finite-size effects.

### B. Jump-driven growth and the core-halo structure of mutant clones

Some typical outcomes of the simulation model are shown in Fig. 1 for both two-dimensional (2D) and onedimensional (1D) ranges. To emphasize variations in the spatial patterns for the same average clone size, simulations were chosen in which the final state has exactly ten unique alleles; this required varying the rescaled mutation rate as *μ* was increased. This feature, which is tied to the slower growth of individual clones apparent in the space-time plots of Fig. 1(b), is explored in depth in Section II.C.

**FIG. 1.**
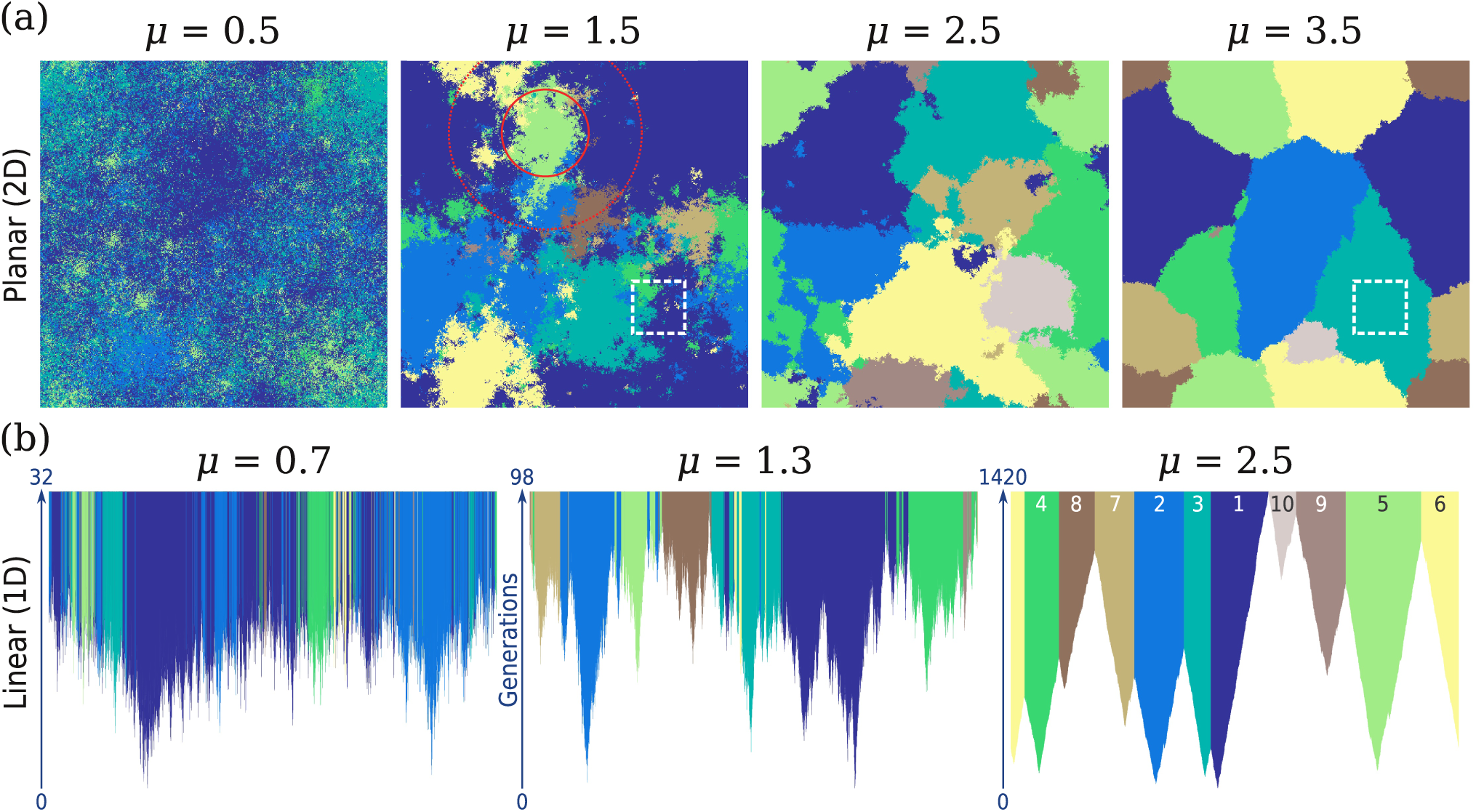
Spatial soft sweep patterns with the same number of distinct alleles vary strongly with jump kernel. (a). Final states of 2D simulations on a lattice of size 512 × 512, for a range of values of the kernel exponent *μ*. Each pixel corresponds to a deme, and is coloured according to the identity of the allele occupying that deme; demes belonging to the same mutant clone share the same colour. Simulations were chosen to have ten unique alleles in the final state; colours reflect the temporal order of the originating mutations as labeled in the lower right panel. Rescaled mutation rates are 3×10^−6^,10^−6^, 10^−6^, and 10^−7^ for kernel exponents 0.4, 1.5, 2.5, and 3.5 respectively. The solid and dotted circles in the second panel indicate the extent of the core and the halo respectively for the light green clone, as quantified by the measures *r*_eq_ and *r*_max_ respectively (see Table II). The subrange highlighted by a dashed box contains six distinct alleles for *μ* = 1.5 but only one allele for *μ* = 3.5. (b) Full time-evolution of 1D simulations with *L* = 16384 for three kernel exponents, chosen so that the final state has ten unique alleles. Each vertical slice displays the lattice state at a particular time (measured in generations), starting from an empty lattice (white) and continuing until all sites are filled and the sweep is completed. The rescaled mutation rates are 3 × 10^−5^, 7 × 10^−6^, and 7 × 10^−7^ respectively from left to right. In the last panel, the colours are labelled according to the order of appearance of the originating mutation; the same order is shared among all panels in the figure.

In both 2D and 1D, the spatial soft sweep patterns of Fig. 1 display systematic differences as the kernel exponent is varied. Clones are increasingly fragmented as the kernel exponent is reduced; i.e. as long-range dispersal becomes more prominent. At the highest value of *μ* in each dimension, the range is divided into compact, essentially contiguous domains each of which shares a unique mutational origin. As the kernel exponent *μ* is reduced, the contiguous structure of clones is lost as they break up into disconnected clusters of demes. For most clones, however, a compact region can still be identified in the range which is dominated by that clone (i.e. the particular allele reaches a high occupancy that is roughly uniform within the region but begins to fall with distance outside it) and in turn contains a significant fraction of the clone. We call this region the *core* of the clone. The remainder of the clone is distributed among many satellite clusters which produce local regions of high occupancy for a particular clone. The satellites become increasingly sparse and smaller in size as we move away from the core. For the broadest kernels (*μ* = 0.5 in 2D and *μ* = 0.7 in 1D), most clones also include isolated demes which do not form a cluster but are embedded within cores and satellite clusters of a different allele. We term the collection of satellites and isolated demes the *halo* region surrounding the core of the clone. The circles in the second panel of Fig. 1(a) illustrate the extent of core and halo, quantified via distance measures which we introduce later on (see Table II) for a particular clone (the fifth clone entering the population, colored light green). The spatial extent of the clone including the halo can be many times the extent of the core alone, and increases relative to the core extent as *μ* is reduced. (We will use “extent” to refer to linear dimensions, and “mass” or “size” to refer to the number of demes.)

The space-time evolution displayed in Fig. 1(b) for linear simulations reveals the role of jump-driven growth in producing the observed spatial structures. At *μ* = 2.5, the growth of clones appears nearly deterministic, with fronts separating mutant from wildtype advancing outwards from the originating mutations at near-constant velocity. These fronts are arrested when they encounter advancing fronts of other clones, leaving behind a tessellation of the range into contiguous clones. By contrast, at the lower values *μ* = 1.3 and 0.7, the stochastic nature of jump-driven growth becomes apparent. Clones advance through long-distance dispersal events, which seed satellite clusters that may merge with each other before the sweep is complete. For all except the smallest clones, the originating mutation is surrounded by a region which is dominated by that particular allele—these form the core regions defined above. Satellites are seeded by stochastic jumps that extend over regions which either were occupied by a different allele already, or get filled in by a different allele before the satellite has a chance to merge with the core. For *μ* = 1.3, haloes extend only a short distance out from the core, whereas at *μ* = 0.7 the haloes often extend over a distance many times the core extent.

The increased fragmentation of clones with broader dispersal kernels has a marked impact on local diversity in sub-regions of the range. Haloes belonging to different alleles overlap to produce regions of high diversity, as exemplified by the dashed box in Fig. 1(a) for *μ* = 1.5, which contains demes belonging to six of the 10 unique alleles despite being a small fraction of the total range area. By contrast, the same region contains only one allele at *μ* = 3.5 for which clones form contiguous domains. Other effects of broadening the dispersal kernel are also visible in Fig. 1: the spread in clone sizes becomes larger, and individual clones take many more generations to attain a given size.

To build a quantitative understanding of these variations, we begin by noting that at early times in Fig. 1(b), each clone grow largely unencumbered by other clones. We can therefore gain insight from existing results on the jump-driven growth of a *solitary* advantageous clone expanding into a wildtype background [17]. The key features are summarized here and illustrated for the blue clone in Fig. 2. Consider a clone that grows from a mutation that originated at time *t* = 0 at the origin. At times longer than a short transient, the clone fills most sites out to some distance from the origin. In line with the terminology established above, we call this region of high occupancy the *core* of the growing clone. Its typical extent over time (i.e. the average radius of a core that has grown for time *t*) is quantified by a function *ℓ*(*t*) which itself depends on the dispersal kernel (a precise definition is given at the end of this section). As sites in the core get filled, they send out offspring through long-range dispersal events drawn from the specified kernel, which then grow into independent satellite clusters. As a result, at any time *t* there are also demes outside the core which are occupied by the mutant. However, the occupancy of sites outside the core decays as *r*^−(*d*+*μ*)^ with distance *r* from the originating mutation [17], fast enough that the total mass of the clone at time *t* is proportional to *ℓ*^*d*^(*t*).

**FIG. 2.**
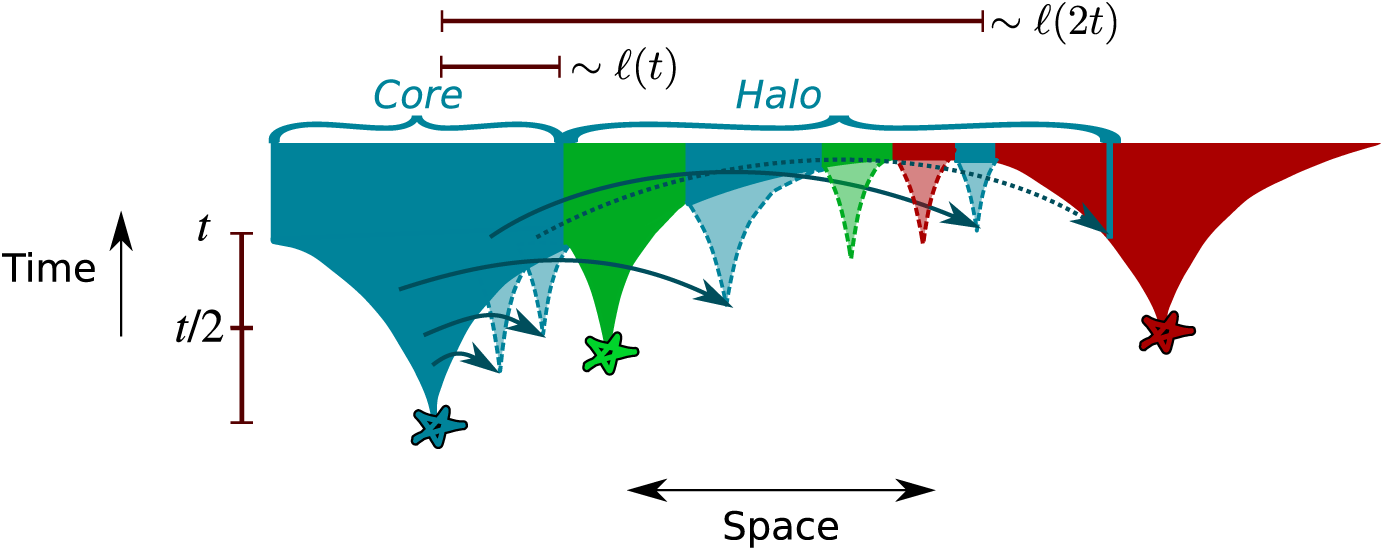
Marginal dynamics of jump-driven soft sweeps. When long-range dispersal is significant, clones of different allelic identity (distinguished by colors) grow out of their originating mutations (stars) by accumulating satellite clusters (translucent cones). If only a single mutation were present, the extent *ℓ*(*t*) of the high-occupancy core (consisting of satellite clusters which have merged with the cluster growing out of the originating mutation) would, on average, follow a faster-than-linear growth rule, depicted schematically by the border of the opaque regions. For a wide range of kernels in the vicinity of *μ* = *d*, the growth rule arises from a hierarchy of length and time scales related by doublings in time: the satellite that merges with the core at time *t* would typically originate from a key jump out of the core at time *t*/2, that extended over a length of order *ℓ*(*t*) (solid arrows out of blue region). Looking forward in time, a clone whose core has grown for time *t* without being obstructed will have likely seeded satellites out to a distance of order *ℓ*(2*t*). In the presence of recurrent mutations, these satellites may be obstructed from merging with the core due to intervening cores and satellites of different mutational origin (green and red regions). For very broad dispersal kernels, the halo also includes rarer jumps out of the core (dashed arrow) that land in regions that are being taken over by other alleles, but establish themselves in stochastic gaps in those regions. The disconnected satellite clusters and isolated demes comprise the halo region of the mutant clone.

As sketched in Fig. 2, the core grows through mergers of satellite clusters that grew out of rare but consequential “key jumps” out of the core at earlier times (solid arrows in Fig. 2). Ref. 17 identified qualitative differences in the behaviour of key jumps and the resulting functional forms of *ℓ*(*t*) as the kernel exponent is varied. When *μ* > *d* + 1, the extent of typical key jumps remains constant over time, which implies that they must originate and land within a fixed distance from the boundary of the high-occupancy region at all times. As a result, clones advance via a constant-speed front similar to the case of wavelike growth; i.e. *ℓ*(*t*) ∝ *t*. Furthermore, the separation between the core and satellites is insignificant at long times, giving rise to essentially contiguous clones. By contrast, for *μ* < *d* + 1, growth is increasingly driven by jumps that originated in the interior of the core at earlier times, and key jumps become longer with time. The resulting growth of *ℓ*(*t*) is faster-than-linear with time. The value *μ* = *d* is an important marginal case which separates two distinct types of long-time asymptotic behaviour for *ℓ*(*t*): power-law growth for *d* < *μ* < *d* + 1 and stretched-exponential growth for 0 < *μ* < *d* (see the second column of Table I for the asymptotic growth forms in all regimes). As *μ* ⟶ 0, spatial structure becomes increasingly irrelevant and the growth dynamics approaches the exponential growth of a well-mixed population.

**TABLE 1.** Core growth asymptotics and characteristic core sizes. The table catalogues the asymptotic behaviour of *ℓ*(*t*) (from Ref. 17) along with the expected scaling of the characteristic clone size *χ*_*as*_, omitting distance and time scales for *ℓ* and *t* respectively. *W* is the Lambert *W*-function, *B*_*μ*_ *≈* 2*d* log(2)*/*(*μ – d*)^2^, and *η* = log[2*d/*(*d* + *μ*)]*/* log 2. The subscript in *χ*_as_ indicates that the asymptotic *ℓ*(*t*) was used as opposed to the more accurate functions listed in Appendix A. Note that for values of *μ* near *d*, the asymptotic growth forms are of limited value since the time befor the asymptotics is reached becomes very large. In this situation, *χ* must be computed using the more accurate scaling forms, see Appendix A for more details.

| Kernel exponent | Asymptotic $\ell(t)$ behaviour | $\chi_{\text{as}}$ |
| --- | --- | --- |
| $\mu < d$ | $\exp(B_\mu t^\eta)$ | $[\xi/W(\xi)]^{\frac{1}{\eta}}, \quad \xi \equiv B_\mu \eta d (\tilde{u} \omega_d)^{-\eta}$ |
| $\mu = d$ | $\exp(c \log^2 t)$ | $(\tilde{u} \omega_d)^{-1} e^{2(\log 2 - \sqrt{\log 2 \log(2/\tilde{u} \omega_d)})}$ |
| $d < \mu < d + 1$ | $t^{1/(\mu-d)}$ | $(\tilde{u} \omega_d)^{-1/\mu}$ |
| $\mu = d + 1$ | $t \log t$ (1D) | $\frac{1}{2} \sqrt{\frac{W(2/\tilde{u})}{2\tilde{u}}}$ |
| $\mu > d + 1$ | $t$ | $(\tilde{u} \omega_d)^{-1/(d+1)}$ |

These features of solitary-clone growth can be directly connected to the spatial patterns in Fig. 1 when recurrent mutations are allowed. The tessellation of the range into contiguous domains for the highest values of *μ* is exactly as expected from the wavelike growth situation when *μ > d* + 1. When *μ* < *d* + 1, by contrast, each clone consists of a growing core and well-separated satellite clusters at any time. Unlike the solitary-mutant case, satellites belonging to a particular clone are no longer guaranteed to merge with the core or with each other at later times: due to allelic exclusion, mergers are obstructed by cores and satellites with a different allelic identity, as shown schematically in Fig. 2. The final pattern of frozen-in satellite clusters comprises the previously identified halo structure around each core when *μ* < *d* + 1.

*Notation and definitions:* Before we proceed, we summarize the various quantities in our analysis, and the conventions used in representing them. (A complete list of variables and definitions is provided in Table II.) One set of physical quantities, represented as Latin symbols without a time argument, measures properties of individual clones after the soft sweep has been completed; i.e. quantities measured from the final simulation outputs such as those displayed in Fig. 1. (These quantities could also, in principle, be measurable from a real spatial population that has recently experienced a sweep.) Of these, quantities that have dimensions of length are the mass-equivalent clone radius *r*_eq_ and the clone extent *r*_max_ (defined in Table II). The solid and dotted circles in Fig. 1(a) illustrate these quantities for a specimen clone. The final clone mass is designated by the symbol *X*. Ensemble averages of these quantities for a given set of model parameter values, obtained by averaging first over all clones within a single simulation and then across many independent simulations, are denoted by ⟨…⟩.

**TABLE 2.** Glossary of length and size scales. Table summarizes the various characteristic lengths (denoted by Greek letters) and measured quantities (masses in capital Roman letters and lengths in small Roman letters). The characteristic time *t*\* is implicitly defined in Eq. 1. Except where explicitly noted, the definitions are valid in all dimensions.

| Symbol | Description | Definition |
| --- | --- | --- |
| *Measures of individual clones in final state of spatial sweep* |  |  |
| $X$ | Final clone mass | Number of demes belonging to clone |
| $r_{\text{eq}}$ | Mass-equivalent clone radius | $(X/\omega_d)^{1/d}$ |
| $r_{\text{max}}$ | Clone extent | Half the separation of pair of demes in clone that are furthest apart (1D)<br>$r_{\text{max}}^8 = \text{eighth central moment of clone (2D)}$ |
| *Averages of final-state quantities* |  |  |
| $\langle q \rangle$ | Ensemble average of quantity $q$ | A range average of $q$ is obtained by averaging over all clones in an individual simulation; the range average is itself averaged over multiple independent simulations to obtain $\langle q \rangle$. |
| $q_{\text{ave}}$ | Expected value of $q$ under mutation-expansion balance | $E[q]$ in the limit $L \gg \chi$ |
| *Measures of growth of solitary clones* |  |  |
| $M(t)$ | Time evolution of clone mass | Number of demes colonized at time $t$ starting from a single deme at time 0. |
| $\ell(t)$ | Expected core radius at time $t$ | $E[(M(t)/\omega_d)^{1/d}]$ |
| *Characteristic lengths from mutation-selection balance* |  |  |
| $\chi$ | Characteristic core extent | $\ell(t^*)$ |
| $\psi$ | Characteristic extent of satellite clusters | $\ell(2t^*)$ |
| $\zeta$ | Characteristic outer limit of rare jumps | $\tilde{u}^{-1/\mu}$ |
| $\lambda_{\text{as}}$ | Asymptotic estimate of length scale $\lambda$ | Characteristic length computed using asymptotic forms for $\ell(t)$ from Table I. |

Our analysis connects these properties of the final, static soft sweep pattern to the dynamic growth behaviour of a *solitary* clone under the same dispersal kernel, in the absence of interference from other clones. For a given dispersal kernel, the typical growth behaviour is captured by the core growth function *ℓ*(*t*) which we introduced previously. A precise definition of *ℓ*(*t*) requires making a choice about how to identify the core region. In contrast to the case of wavelike growth, there is no sharp advancing front which separates the high-occupancy region of a growing clone from its surroundings; the average radial occupancy profile at time *t* (defined as the probability that a deme at distance *r* from its point of origin is occupied by the clone) is close to one out to some distance from the origin, beyond which it crosses over to a profile that decays as a power law with increasing distance. One possibility, proposed in Ref. 11, is to define *ℓ*(*t*) as the distance at which the average occupancy profile falls below some low threshold probability *ε*. Here, we make a different choice motivated by the property, proved in Ref. 17, that the total mass of the clone (which we call *M*(*t*)) is proportional to *ℓ*^*d*^(*t*). We define *ℓ*(*t*) as the expected mass-equivalent radius of the clone at time *t*: *ℓ*(*t*) *≡* E[(*M*(*t*)*/ω*_*d*_)^1*/d*^], where *ω*_*d*_ is the volume of the *d*-sphere of radius 1 (*ω*_1_ = 2, *ω*_2_ = *π*). For a particular solitary-clone growth simulation, *M*(*t*) is straightforward to measure since the clone mass is readily accessible. For a particular value of *μ*, *ℓ*(*t*) is then estimated using an ensemble average over many independent solitary-clone simulations (see Appendix A for details). Our choice of *ℓ*(*t*) is proportional to *ℓ*(*t*) defined using an occupancy threshold, provided *ε* is small enough. We expect that using other definitions of to measure since the clone mass is readily accessible. For a particular value of *μ*, *ℓ*(*t*) is then estimated using an ensemble average over many independent solitary-clone simulations (see Appendix A for details). Our choice of *ℓ*(*t*) is proportional to *ℓ*(*t*) defined using an occupancy threshold, provided *ε* is small enough. We expect that using other definitions of *ℓ*(*t*) which scale proportionately with the core region will not significantly change our results, at most shifting the magnitude of reported quantities by constant factors of order unity as long as we are sufficiently far from the well-mixed limit *μ* → 0.

Finally, the interplay between the expansion of individual clones and the introduction of new mutations is used to derive various time-independent characteristic lengths, which are represented as Greek symbols. These length scales depend on the dispersal kernel via the functional form of *ℓ*(*t*), and the rescaled mutation rate 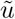. Precise definitions of the characteristic length scales are provided in Table II and in the forthcoming sections.

#### 1. Marginal dynamics and the relative sizes of core and halo

We can quantify the expected spatial extent of entire clones (including haloes) relative to cores by considering the dynamics in the vicinity of the marginal value *μ* = *d*. Although the long-time asymptotic dynamics are qualitatively different above and below this value (power-law in *t* for *d* < *μ* < *d* + 1, and stretched-exponential for 0 *< μ < d*), the approach to the asymptotic behaviour is extremely slow for values of *μ* close to *d*, with the intermediate-time evolution controlled by the marginal dynamics at *μ* = *d*. As a result, the marginal dynamics is important for a wide range of values of *μ* at biologically-relevant time scales [17].

In the marginal regime, the scaling behaviour of key jumps follows a particularly simple pattern, illustrated schematically in Fig. 2: satellite clusters which merge with the core at time *t* are seeded by key jumps that typically happened around time *t*/2 and covered a distance of order *ℓ*(*t*) ≫ *ℓ*(*t*/2). Therefore, a core that has grown up to some extent of order *ℓ*(*t*) has likely already seeded satellites out to a distance of order *ℓ*(2*t*), some of which will have reached an appreciable size as illustrated in Fig. 2. If the core has grown to some linear size *l*, we then expect satellites that have reached a significant size to extend to a distance *l*^*′*^ = *ℓ*(2*ℓ*^−1^(*l*)), which may be considered to be a lower bound on the expected extent of the halo. When isolated demes embedded within cores and satellites belonging to other clones are included, the full spatial extent of the clone is even larger, because there remains a finite probability of rare jumps from the core out to distances farther than *l*^*′*^ (dotted arrow in Fig. 2).

The above estimate for *l*^*′*^, when approximated using the long-time asymptotic growth rules for different jump kernels, reveals qualitatively different scaling behaviours for the clone extent on either side of the critical point *μ* = *d*. For power-law growth, *ℓ*(*t*) *∼ t*^1*/*(*μ–d*)^, we find *l*^*′*^*/l ∼* 2^1*/*(*μ–d*)^; i.e. the ratio of halo size to core size is a constant that grows as *μ → d* but is independent of the size of the clone. By contrast, in the stretched-exponential regime, with *ℓ*(*t*) *∼* exp(*Bt*^*η*^) where *B* and *η* depend on *μ*, we find 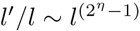; i.e. the ratio of halo to core depends on the core size as well as on the kernel exponent. Since *η* = log[2*d/*(*d* + *μ*)]*/* log 2 *>* 1, the halo becomes increasingly prominent as *μ* → 0. These scaling estimates break down as *μ* approaches *d*—for instance, the ratio *l*^*′*^*/l* diverges in the power-law growth regime—mirroring the limited utility of the approximate asymptotic forms for *ℓ*(*t*) near *μ* = *d*. Instead, we must use the more accurate forms for *ℓ*(*t*) (Appendix A) to evaluate *l* and *l*^*′*^. However, upon using these forms with fitted magnitude and time scales, the qualitative picture is largely unchanged: the ratio *l*^*′*^*/l* becomes very weakly dependent on the core size as *μ* ↘ *d*, but the dependence is much stronger when *μ* drops below *d*. For instance, at *μ* = 1.2 in 1D, the predicted ratio *l*^*′*^*/l* merely doubles from 4 to 8 as *l* spans four orders of magnitude from *l* = 10 to *l* = 10^5^; by contrast, at *μ* = 0.8 the ratio changes by an order of magnitude (from roughly 5 to 70) over the same range of core sizes.

In summary, the growth dynamics for solitary clones at different kernel exponents predicts the following structure for large mutant clones: contiguous, compact clones for *μ > d* + 1; a high-occupancy core with a halo of well-developed satellite clusters that extends out to a size-independent (but kernel-dependent) multiple of the core radius for *d* < *μ* < *d*+1; and a sparse halo which is significantly larger in extent than the core and becomes more prominent with increasing clone size for *μ < d*. We now assume that these conclusions, and the associated scaling relations for the linear extent of the core and halo, also apply to the spatial structure of mutant clones that have grown during soft sweeps and have been frozen in due to interference with clones of differing mutational origin. This key assumption is tested in the following section.

#### 2. Occupancy profiles

To verify the structural features outlined above, we measured average occupancy profiles of distinct clones in the final states of 1D soft sweep simulations (Fig. A1). Occupancy profiles from clones of different sizes are combined by scaling the distance coordinate of each profile by the mass-equivalent radius *r*_eq_, derived from the total mass *X* of that clone via *r*_eq_ *≡* (*X/ω*_*d*_)^1*/d*^, and performing an ensemble average as described in Table II. The choice of distance scale is motivated by our definition of *ℓ* in terms of the clone mass, and justified by the observation that averaged occupancy profiles for a given kernel with vastly different average clone sizes collapse onto a single curve when the distance coordinate is rescaled by the size, consistent with the core radius being proportional to *r*_eq_, see Supplementary Fig. A1.

Ensemble-averaged occupancy profiles for different jump kernels are shown in Fig. 3(a) with 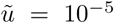. We observe that when *μ > d* + 1, the averaged occupancy is negligible for *r/r*_eq_ *>* 2, and the curve has a point of symmetry at (*r/r*_eq_, ⟨*ρ*⟩) = (1, 1*/*2), such that ⟨*ρ*⟩(*r/r*_eq_) = 1 −⟨*ρ*⟩ (2 *r/r*_eq_) for 1 *−< r/r*_eq_ *<* 2. This form is consistent with the entire clone being contained in a single domain, which grows to different lengths on either side of the originating mutation, as illustrated in the inset.

**FIG. 3.**
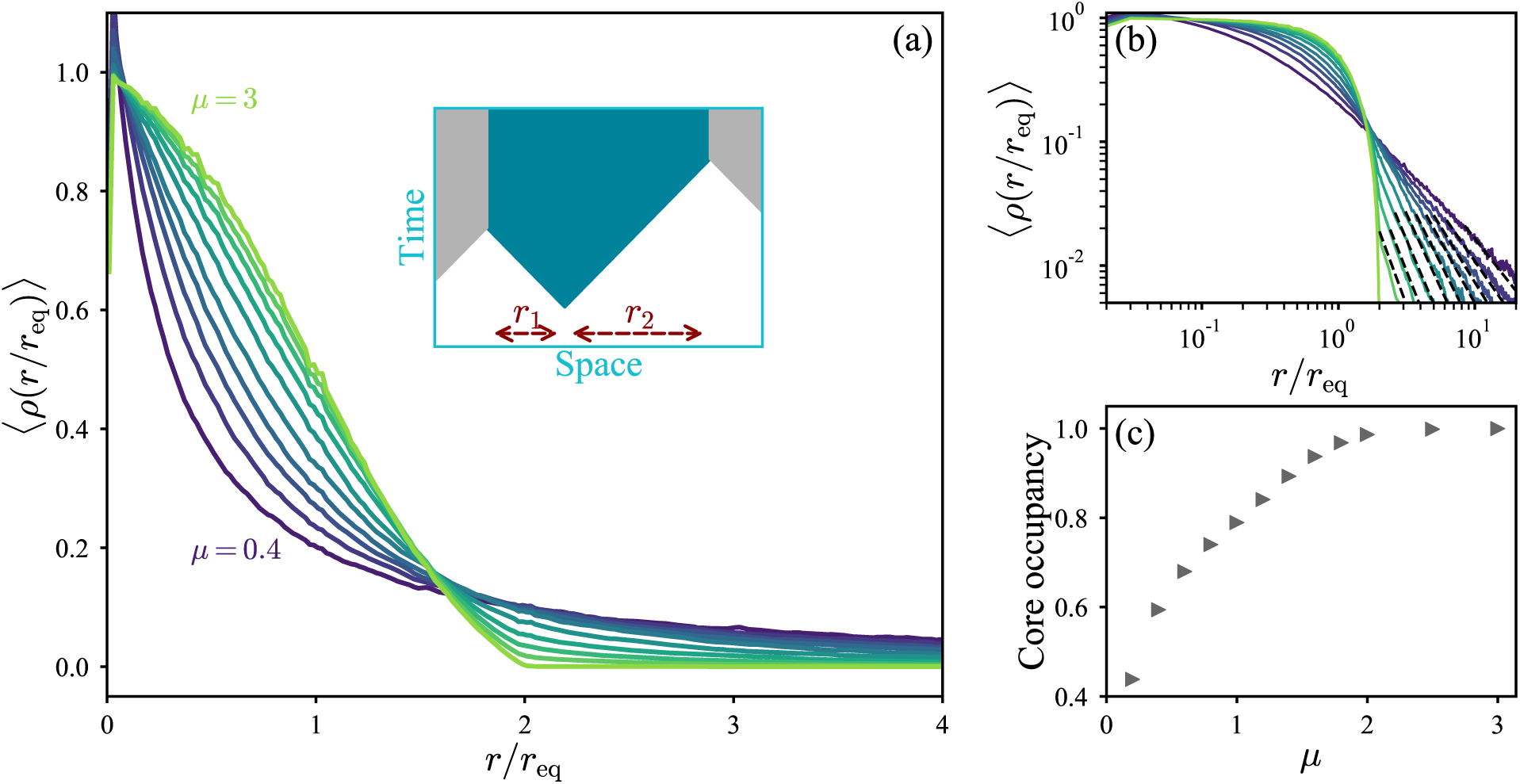
Occupancy profiles of clones show the reduced prominence of the core with increased long-range dispersal. (a) Ensemble-averaged occupancy profiles of mutant clones in 1D, with *L* = 10^6^ and 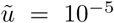. The occupancy profile of a particular clone is defined as the probability *ρ*(*r*) that a deme at distance *r* from its point of origin is occupied by that clone. Colours signify different dispersal kernels, with exponents *μ* = {0.4, 0.6, 0.8, 1, 1.2, 1.4, 1.6, 1.8, 2, 3} in order of increasing occupancy at the origin. Curves are obtained by averaging individual occupancy profiles from all clones with total mass *X >* 100 to obtain a range-averaged occupancy profile ⟨*ρ*⟩ for each of 100 independent simulations for each dispersal kernel; these were then averaged to obtain the ensemble-averaged profile 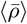. Inset illustrates the origin of the variation in occupancy for *r/r*_*eq*_ *<* 2 in the wavelike growth regime: the mutational origin need not be positioned at the centre of mass of the contiguous domain, giving rise to a single-clone occupancy profile *ρ*(*r*) = {1, 0 *< r* ≤ *r*_1_; 1*/*2*, r*_1_ *< r* ≤ *r*_2_; 0*, r > r*_2_}. (b) Same data as in (a) on logarithmic scales. The dashed lines show the power-law dependence ⟨*ρ*⟩ *∝ r*^−(*μ*+*d*)^. (c) Core occupancy, defined as the fraction of the ensemble-averaged occupancy that lies within 0 *< r/r*_*eq*_ *<* 2, as a function of kernel exponent *μ*.

The predicted breakup of clones due to long-range dispersal is reflected in the overall broadening of occupancy profiles as the kernel exponent *μ* is reduced below the critical value *d* + 1. In this range, an appreciable portion of the clone lies outside the maximal distance from the origin (*r/r*_eq_ = 2) that could be measured for a contiguous domain in a linear habitat. (This upper limit would correspond to a clone that was obstructed by an occupied deme adjacent to its mutational origin on one side, and attained its final mass by expanding only in the other direction.) The dropoff in occupancy becomes increasingly steep at low values of *r/r*_eq_ as *μ* is reduced, but more gradual at larger distances, consistent with a narrowing high-occupancy region balanced by a halo of increasing prominence. At large distances, the falloff in occupancy is consistent with the power-law behaviour expected from the solitary-clone growth dynamics, ⟨*ρ*⟩ *∝ r*^−(*μ*+*d*)^ [dashed lines in Fig. 3(b)], which supports our assumption that the final structure of a mutant clone in a spatial soft sweep is similar to that of a solitary clone expanding without interference.

To quantify the relative prominence of the halo to the core across all growth regimes, we define the core occupancy as the fraction of the total occupancy contained within the maximal range of distances that could be measured for contiguous domains, 0 *< r/r*_eq_ *<* 2. We find that the core occupancy is close to 100% for *μ > d* + 1 (= 2 in 1D), consistent with the contiguous clones and insignificant haloes expected for wavelike growth [Fig. 3(c)]. For broader kernels, the core occupancy falls with *μ*, reflecting the increasing prominence of the halo as *μ* approaches zero. However, the core still contains an appreciable fraction of the total occupancy for all values of *μ* that we have simulated. This observation further supports our approach of connecting the geometric extent of cores to the total mass of their corresponding clones, as we do in the following section.

### C. Characteristic scales via mutation-expansion balance

So far, we have focused on the spatial structure of individual clones within a soft sweep, and have shown that many aspects of this structure can be understood from the theory of growth of a solitary clone under the same dispersal kernel. To address questions of global and local allelic diversity, however, we need to explicitly consider the concurrent growth of multiple clones. We now show how the balance between jump-driven growth and the dynamics of introduction of new mutations sets the typical size and spatial extent of clones.

#### 1. Size of a “typical” clone

In an infinitely large range, a solitary clone could grow without bound, but in the presence of recurrent mutations, the growth of any one clone is obstructed by other clones. Balancing mutation and growth gives rise to a characteristic time scale *t*\*, and associated characteristic linear extent *χ*, for mutant clones in multi-origin spatial sweeps [11]. These scales determine whether a sweep will be “hard” or “soft” within a finite range of given size.

When clones grow as compact, connected domains, growth is interrupted when the advancing sharp boundary of the clone encounters a different allele. However, for clones growing via long-range dispersal events, a sharp boundary no longer exists, and small obstacles can be traversed by jumps. The picture of jump-driven growth that we have developed suggests that haloes belonging to different clones can freely overlap, whereas core regions cannot. Therefore, new mutations arising within the halo region of a growing clone do not significantly impede its growth. Instead, the crucial factor restricting the growth of a clone is when its high-occupancy *core* encounters a different clone, as depicted schematically in Fig. 2. Since *ℓ*(*t*) defines precisely the time-evolution of the core extent of a solitary clone, we define *t*\* as the time interval for which exactly one mutation is expected to occur in the space-time region swept out by the growing core:

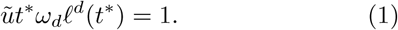

The corresponding characteristic extent, *χ =ℓ*(*t*\*), matches the length scale introduced in Ref. 11 to characterize spatial soft sweeps.

Rough estimates for *t*^*\**^ and *χ* can be obtained by using the long-time asymptotic forms for *ℓ*(*t*) in the different growth regimes, see Table I. These estimates highlight the vastly different functional dependences of the characteristic scales on the rescaled mutation rate as the kernel exponent is varied. For quantitative tests, we use scaling forms for *ℓ*(*t*) derived in Ref. 17, which are much more accurate at short times when *µ* is close to *d*. All theoretical forms for *ℓ*(*t*) include unspecified multiplicative factors *A* and *B* for the length and time variables, which are fixed by fitting the functional forms to the growth of isolated clones starting from a solitary seed, see Appendix A for details.

The scales *χ* and *t*^*\**^ provide the appropriate rescaling of space and time to compare two sweeps with different mutation rates but the same growth rule *ℓ*(*t*), and therefore capture the dependence of many soft sweep features on the mutation rate. Most significantly, they set the expected number of independent mutational origins in a range of a given size. When both sides of Eq. 1 are multiplied by the total number of demes *L*^*d*^, it equates to a condition for the range to be completely filled by mutations accumulated at a rate 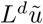 over a time *t*^*\**^ without interference, each of which grows to the characteristic size *ℓ*(*t*^*\**^). The expected number of mutational origins in the range therefore scales as 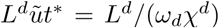. If 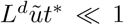, or equivalently *L « χ*, it is unlikely that many independent mutations will arise: the sweep is likely to be hard. By contrast, if the range is large compared to the characteristic length *χ*, the number of independent origins grows in proportion to the range area. Consequently, the total number of demes in the range divided by the expected number of origins converges to a well-defined value as *L* increases, which we call the expected clone mass *X*_ave_. For a given dispersal kernel, the mutation-rate dependence of *X*_ave_ is captured by the variation of *χ* with 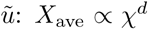, with a 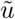-independent factor of proportionality that dependes on the details of the growth dynamics.

To test whether the characteristic length scale quantifies the number of mutational origins in a range, we compare the ensemble-averaged clone mass measured in simulations, ⟨*X*⟩, to 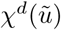 computed using the theoretical forms for *ℓ*(*t*). Results for 1D simulations are shown in Fig. 4. (Definitions of measured quantities and averages are provided in Table II.) For clarity, the expected scaling with mutation rate in the wavelike spreading limit, 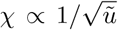, is divided out. We find that the average clone sizes for different system sizes coincide at 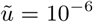, consistent with ⟨*X*⟩ being an estimate of an underlying expected clone mass, *X*_ave_, that is determined by the mutation-expansion balance and is independent of system size. For each value of *µ*, the computed 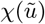 quantitatively captures the dependence of ⟨*X*⟩ on 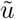 over many orders of magnitude, up to a 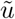-independent constant factor (the factor depends on model details and is treated as a fitting parameter, but it turns out to not vary significantly with *µ*). These results confirm that the mutation-expansion balance captured in Eq. 1, first identified in Ref. 11, remains relevant for characterizing the compact core regions of clones in when long-range dispersal is prominent.

**FIG. 4.**
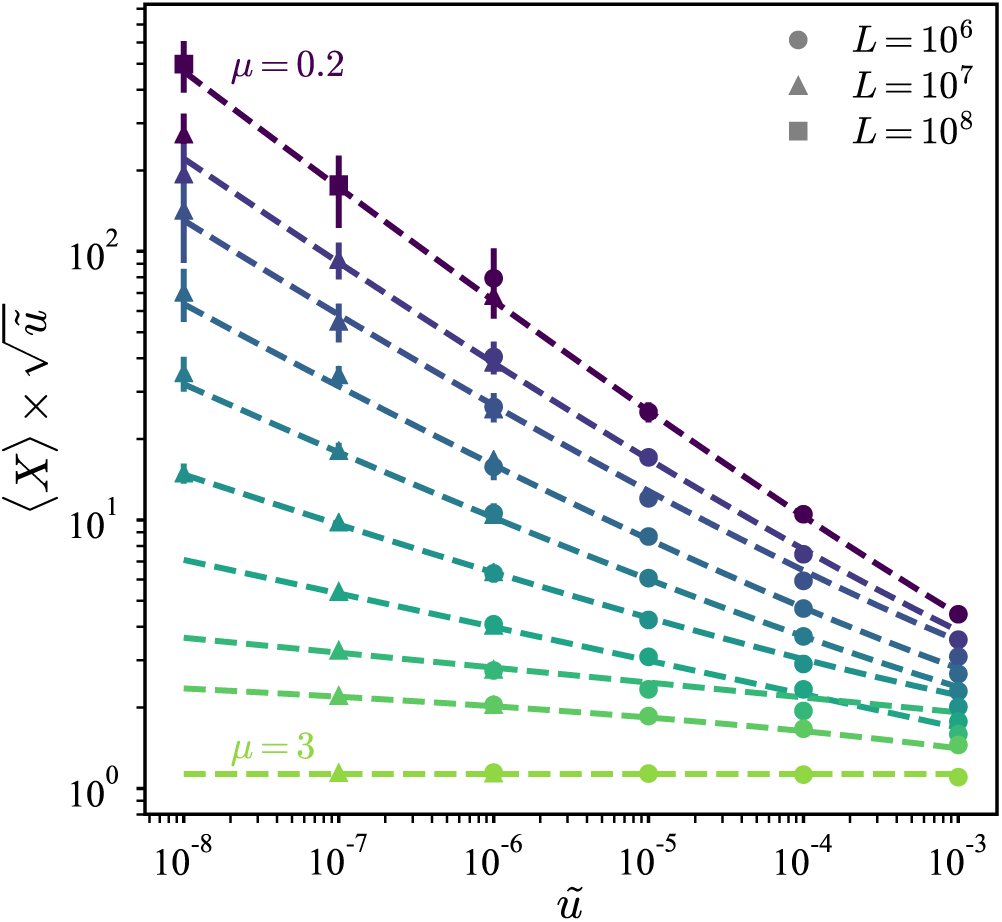
Average clone mass is set by mutationexpansion balance. The ensemble-averaged final mass of mutant clones ⟨*X*⟩ as measured from 1D simulations as a function of rescaled mutation rate, scaled by the expected dependence 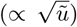 for wavelike growth of clones. Results from different system sizes (symbols) are presented for each dispersal kernel quantified by *µ* (colours); the values are (from top to bottom) *µ* = {0.2, 0.6, 0.8, 1, 1.2, 1.4, 1.6, 1.8, 2, 3 } Each point represents an average over 20 or more independent simulations. Error bars denote measured standard deviation across repetitions. Dashed lines show the theoretical predictions for 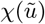 for each dispersal kernel (see Appendix A for details), multiplied by a *µ*-dependent magnitude factor whose value is 1.5 for 1 *≤ µ <* 2, 1.6 for *µ >* 2, and 1.65 for all other values.

#### 2. Characteristic extent of clones

The analysis of the spatial structure of individual clones (Section II.B) showed that the extent of clones (i.e. the portion of the range over which demes belonging to the clone can be found) can be many times larger than the expected extent for a compact clone, especially for broad dispersal kernels. Therefore, the average clone size may not be representative of the spatial extent of typical clones over the range, pointing to the potential relevance of additional length scales when characterizing jump-driven spatial soft sweeps. To quantify this effect, we measure the extent, *r*_max_, of a clone in our 1D simulations as half the largest distance between any pair of individuals belonging to that clone. The disparity between the true extent of clones and the extent expected from the average clone mass can be evaluated by comparing the ensemble-averaged extent ⟨*r*_max_⟩ to the average mass-equivalent radius ⟨*r*_eq_⟩. If all clones were perfectly contiguous and compact, we would expect ⟨*r*_max_⟩ = ⟨*r*_eq_⟩.

Figure 5 shows the ratio of average clone extent to average mass-equivalent radius for different dispersal kernels and mutation rates. The ratio is unity when *µ > d* + 1, which is the expected regime of compact clones. For broader kernels, we find that the average spatial extent is larger than the mass-equivalent radius, consistent with our expectation from jump-driven growth. Two separate behaviours can be identified in this regime. In the range *d < µ < d*+1, the ratio ⟨*r*_max_⟩/⟨*r*_eq_⟩ is independent of the rescaled mutation rate. By contrast, for *µ ≤ d*, the ratio of lengths shows a mutation-rate dependence, and grows dramatically in magnitude. At the smallest values of *µ*, the measured extent of the largest clones becomes limited by the system size (the maximum measurable extent under periodic boundary conditions is *L/*2). This finite-size effect artificially suppresses the ratio ⟨*r*_max_⟩/⟨*r*_eq_⟩, as is apparent upon comparing the measurements at *L* = 10^6^ and *L* = 10^7^. (Note that the measurement of ⟨*r*_eq_⟩ does not suffer from finite-size effects, which was verified in Fig. 4, since the core extent remains much smaller than the system size for these parameters.)

**FIG. 5.**
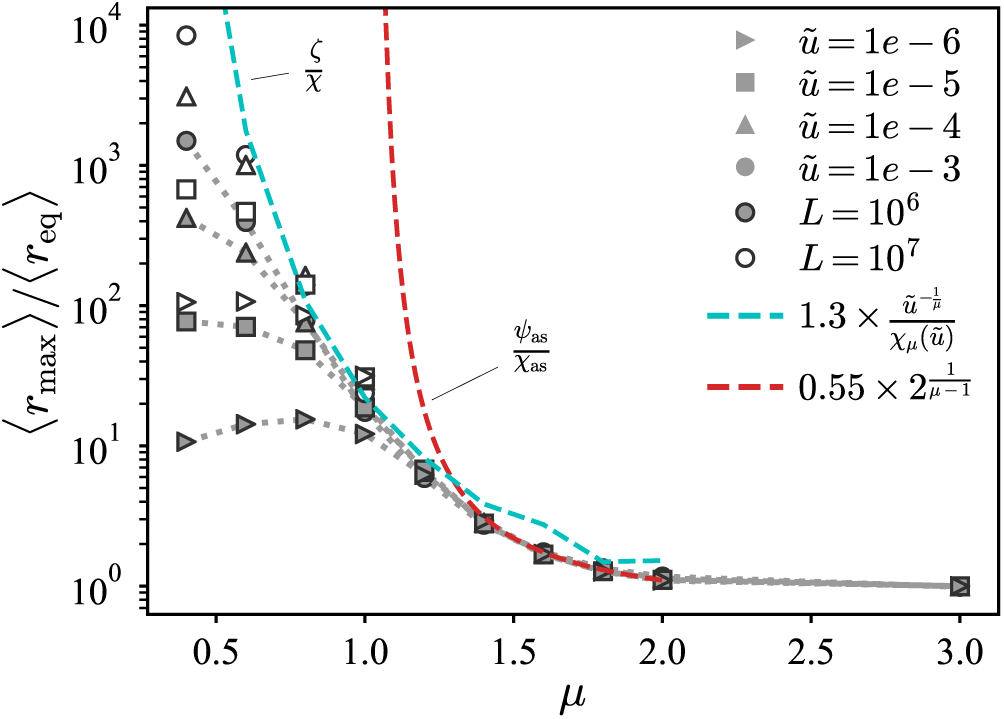
The spatial extent of clones can be much larger than the expectation derived from the average mass. Ensemble-averaged spatial extent of clones ⟨*r*_max_⟩, where *r*_max_ is defined as half the distance between leftmost and rightmost demes belonging to a particular clone. Values are normalized by the ensemble-averaged mass-equivalent radius ⟨*r*_eq_⟩ in 1D simulations. Each point is an average over values from 20 independent simulations. Dotted lines connect simulation data points. Dashed lines show the theoretical expectations *ζ/χ* (with 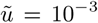) and *ψ*_as_*/χ*_as_ in the ranges *µ <* 1 and 1 *< µ <* 2 respectively, multiplied by model-dependent numerical factors. For *µ <* 1, 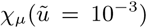 is evaluated as described in Appendix A; for 1 *< µ <* 2, the ratio *ψ*_as_*/χ*_as_ is independent of 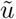. The fall in the measured ratio at low values of 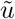 for *µ <* 1 is due to finite-size effects, as seen by comparing the two system sizes: the measured values of *r*_max_ in this range are below the values that would be measured in an infinitely large system.

To explain these features, we return to the core-halo picture of clone structures. We had previously identified two contributions to the halo region of a clone (Fig. 2). One contribution comes from satellite clusters which are established with high probability during the growth process, but are obstructed from merging with the core by clusters belonging to other clones. In addition, the heavy-tailed jump kernel could populate demes arbitrarily far from the growing core. These would form isolated demes or small clusters embedded within other clones, without any related clusters in the neighbourhood. These two mechanisms lead to different characteristic length scales which we now analyze in detail.

We first quantify the extent of the region in which satellite clusters are established. We had argued that, in the vicinity of the critical value *µ* = *d*, a clone whose core has grown to some size *ℓ*(*t*) will have likely established satellites of a significant size out to a distance *ℓ*(2*t*). Having derived a characteristic time scale *t*^*\**^ (defined in Eq. 1) for the growth of a typical clone restricted by mutation-expansion balance, a rough estimate for the extent of its halo is provided by the quantity

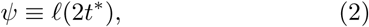

which quantifies the extent of established satellites of the typical clone. Although this estimate ignores interference among haloes of different clones, we note that the expected number of new mutational origins encountered by each satellite cluster is less than one since the satellites are smaller than *ℓ*(*t*^*\**^), which limits the amount of interference. An alternative argument, which balances the rate of key jumps out of the core with the expected rate of mutations arising in the target region of the jumps and therefore incorporates some interference effects, produces a similar estimate for the typical extent of the satellite cluster region (see Appendix C).

Since the existence of satellite clusters is closely tied to the growth process of the core, we expect the typical halo extent to be at least of order *ψ*. In this part of the halo, the distribution of satellite clusters of the same identity as the core is relatively dense: in the absence of interference with other clones, the maximum separation between satellite clusters would be of order *ℓ*(*t*^*\**^) = *χ*. However, the halo also includes contributions from rare long-distance dispersal events which land well outside the dense region of satellite clusters. Due to the heavy tail of the dispersal kernel, the growing core could send out offspring to arbitrarily large distances, which are not restricted by the length scale *ψ*. If these rare jumps land on unoccupied demes, they would establish isolated demes or small clusters. Unlike the satellite clusters, these isolated clusters would be very sparsely distributed, being separated from their relatives by distances much larger than *χ*. However, they would still count as part of the discontiguous halo of their parent core. In particular, the extremal measure *r*_max_ is sensitive to isolated offspring even if they do not belong to satellite clusters of significant size.

The outer limit of jumps made by the core during the sweep can be estimated using prior scaling arguments. Since our time units are set by the migration rate, the net number of jumps out of a typical clone which grows over a time *t*^*\**^ is roughly *ω*_*d*_*t*^*\**^*ℓ*^*d*^(*t*^*\**^), which equates to 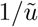 by the definition of *t*^*\**^. The fraction of these jumps which end up beyond a distance *l* is 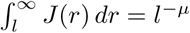. A crude estimate for the outer limit *ζ* of these rare events is obtained by requiring the net number of jumps from the core to distances *ζ* or greater to be equal to 1:

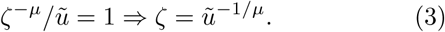

Unlike the extent of the satellite cluster region, *ζ* does not depend explicitly on the core growth function *ℓ*(*t*). However, the scaling for *ζ* does not account for the fact that many of the long-distance rare jumps would fail to establish because they land on the high-occupancy cores of competing clones, which fill up the range over the same time scale *t*^*\**^. Therefore, *ζ* is likely to be relevant when the sparse halo regions become dominant over the cores, i.e. for *µ < d*.

To test whether *ψ* or *ζ* determines the average clone extent in the different growth regimes, we also need to fix an overall magnitude factor, which is not predicted by the scaling arguments. The most general behaviour would be for these magnitude factors to themselves depend on *µ*. However, our measurements of the clone mass (Fig. 4) showed that the magnitude factors only vary slightly over the range of values of *µ*. Therefore, we evaluate the effectiveness of the theoretical length scales by testing whether they reproduce the simulation data up to a *µ*-independent magnitude factor, which we treat as a fit parameter.

The asymptotic growth forms for *ℓ*(*t*) from Table I can be used to obtain the qualitative behaviour of the characteristic satellite cluster extent; we term the resulting estimate *ψ*_as_. (We expect asymptotic estimates to become inaccurate as *µ* approaches *d*.) In the region of power-law growth, *d < µ < d* + 1, we find that *ψ*_as_ and *ζ* both have the same mutation-rate dependence for a given dispersal kernel, 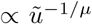. However, the ratios of the length scales to the characteristic core size *χ*_as_ show different behaviour as *µ* is varied: *ψ*_as_*/χ*_as_ = 2^1*/*(*µ–d*)^ has no remaining dependence on *χ*_as_ or *ℓ*(*t*), whereas 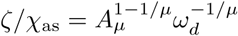 depends on the magnitude parameter *A*_*µ*_ which characterized the solitary-clone growth. The measured ratio of average clone extent to average mass-equivalent radius agrees well with the theoretical prediction for *ψ*_as_*/χ*_as_ for *µ* ≥ 1.4 (red dashed line in Fig. 5; the overall magnitude factor was chosen to match the value at *µ* = 2). By contrast, the prediction *ζ/χ* (cyan dashed line) deviates by factors of order one from the measured ratio due to the residual dependence on *A*_*µ*_, although it agrees with the overall trend. As expected, the asymptotic estimates become increasingly inaccurate as *µ* → 1 which reflects the breakdown of the asymptotic growth form.

At the marginal value *µ* = *d*, the asymptotic form predicts 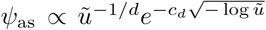 for 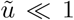, where *c*_*d*_ is a numeric constant of order one. In 1D, the very weak additional dependence on log 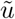 is not sufficient to distinguish this form from the alternative length scale 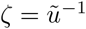 for *µ* = 1. However, the differences in *ψ* and *ζ* become significant in the region of stretched-exponential growth, *µ < d*. As *µ* approaches zero, the strong divergence in Eq. 3 for rescaled mutation rates below one induces *ζ* to grow much faster than *ψ*_as_. For our 1D simulations, the ratio *ψ/χ* only varies by about an order of magnitude, regardless of whether the asymptotic forms or the more accurate scaling forms from Appendix A are used. Therefore, the satellite cluster region cannot account for the dramatic increase in average clone extent observed at low values of *µ* in Fig. 5. By contrast, *ζ* grows rapidly over many orders of magnitude over the same range. Upon fitting an overall magnitude factor, the ratio *ζ/χ*_*µ*_ successfully captures the variation in ⟨*r*_max_⟩/⟨*r*_eq_⟩ for the largest rescaled mutation rate 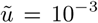 (cyan dashed line in Fig. 5).

To summarize, we have identified two characteristic scales, *ψ* and *ζ* (defined in Eqs. 2 and 3 respectively), that could set the average halo extent in our spatial soft sweeps. Differences between *ψ* and *ζ* are small when *d ≤ µ < d* + 1, but become significant for *µ < d*. Comparisons with the measurements of clone extent in simulations (Fig. 5) support the hypothesis that *ζ* sets the halo extent for the highly sparse clones that arise when *µ < d*, while *ψ* sets the halo extent for the more compact but still discontiguous clones when *µ ≤ d < d* + 1.

### D. Clone size distributions vary with dispersal kernel and influence global sampling statistics

Unlike our simulations, studies of real populations do not have access to complete allelic information over the entire range. Instead, the allelic identity of a small number of individuals is sampled from the population. The likelihood of detecting a soft sweep in such a random sample is determined not only by the total number of distinct clones in the range, but also by their size distribution: if the range contains many clones, but all but one are at extremely small frequency (defined as the fraction of demes in the range that belong to that clone), the sweep is likely to appear “hard” in a small random sample which would with high probability contain only the majority allele. Long-range dispersal can therefore influence soft sweep detection not only by setting the average clone size, but also by modifying the distribution of clone sizes around the average. Having already established that the dispersal kernel has a significant effect on the average clone size (Fig. 4), we now analyze its effects on the clone size distribution and the consequences for soft sweep detection.

Clone size distributions were quantified by computing the *allele frequency spectrum f*(*x*), defined such that *f*(*x*) *δx* is the expected number of alleles which have attained frequencies between *x* and *x* + *δx* in the population [18]. The allele frequency spectrum is related to the average probability distribution of clone sizes, but has a different normalization 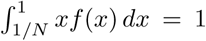 which allows sampling statistics to be expressed as integrals involving *f*(*x*) (we will exploit this fact in Section II.D.1 below). Analytical results for *f*(*x*) can be derived for the deterministic wavelike growth limit *µ » d* + 1 in 1D by mapping the spatial soft sweep on to a grain growth model [19], and for the panmictic limit *µ* → 0 in any dimension via a different mapping to an urn model [20]. The resulting functions, termed *f*_w_ and *f*_*∞*_ for the two limits respectively, provide bounds on the expected frequency spectra at intermediate *µ*. Details of the mappings and complete forms for the functions *f*_w_ and *f*_*∞*_are provided in Appendix D.

Fig. 6(a) shows allele frequency spectra computed from the outcomes of 1D soft sweep simulations for system size *L* = 10^7^ and mutation rate 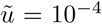 We find that the frequency spectra vary strongly with the dispersal kernel, and approach the exact forms *f*_*∞*_ and *f*_w_ for small and large *µ* respectively. Generically, spectra become broader as the kernel exponent is reduced: as *µ* → 0, more high-frequency clones are observed. Although this broadening is partly explained by the increase in the average clone size due to accelerated expansion, which would lead to more high-frequency alleles, there are also systematic changes in the overall shapes of the distribution as the dispersal kernel is varied. Upon reducing the rescaled mutation rate to 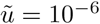 [Fig. 6(b)], all frequency spectra broaden due to the increase in the average clone size, but the variations in shapes of the *f*(*x*) curves with *µ* remain consistent across the two mutation rates. These observations suggest that spatial soft sweep patterns with similar numbers of distinct alleles in a range might nevertheless have vastly different clone size distributions due to different dispersal kernels, with implications for sampling statistics.

**FIG. 6.**
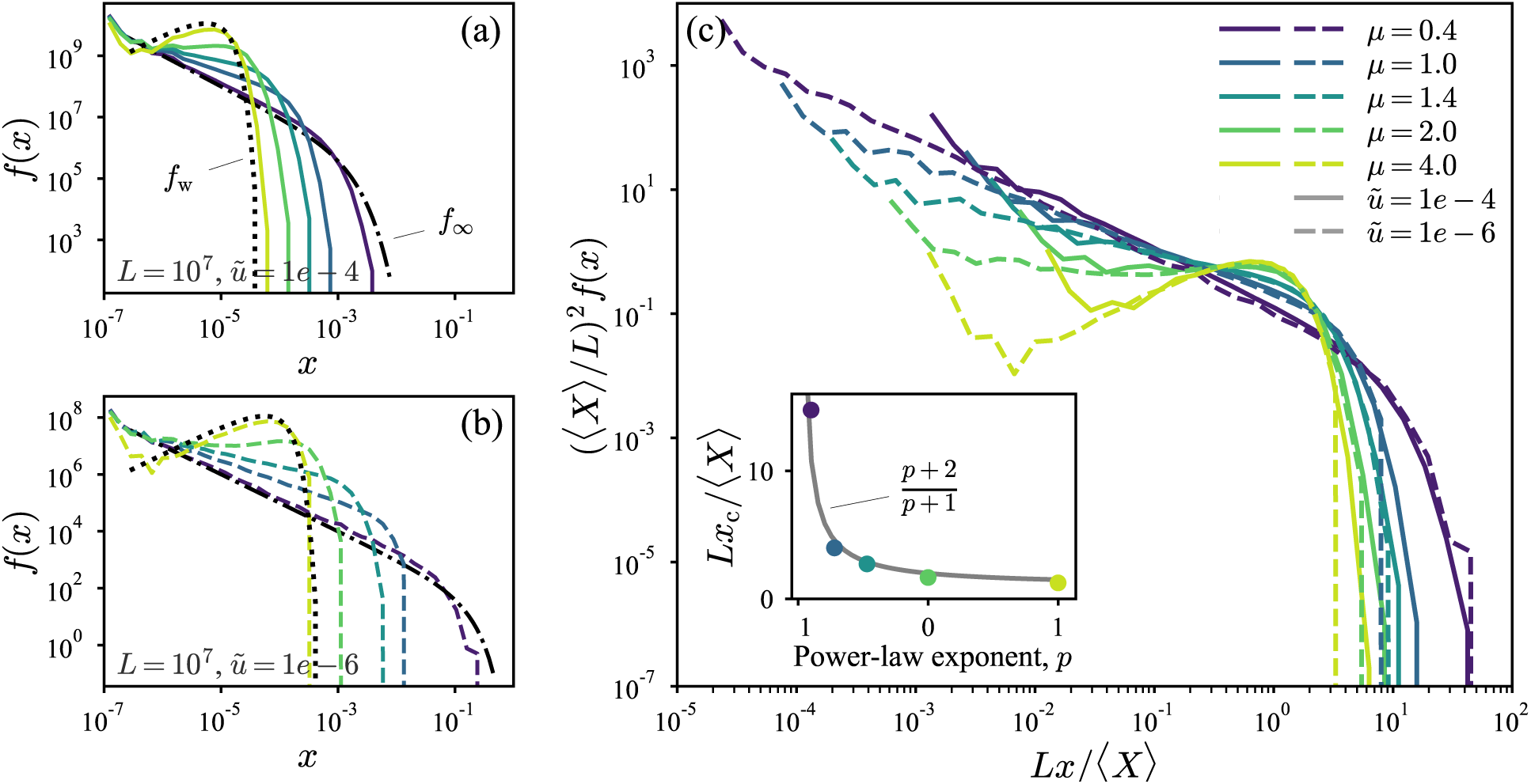
Variation of allele frequency spectra with mutation rate and dispersal kernel. (a) Allele frequency spectra *f*(*x*) estimated from 1D simulations with *L* = 10^7^ and 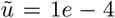. Each curve is the average of frequencies measured from 20 independent simulations. Curves are coloured by dispersal kernel according to the legend in (c). The uptick for the lowest bin is an artifact of the logarithmic bin sizes together with the hard lower cutoff in allele frequency at *x* = 1*/L*. Black dotted and dash-dotted lines show the analytical distributions for wavelike spreading and panmictic limits respectively. (b) Same as in (a) except with 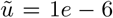 (c) Frequency spectra from (a) and (b), rescaled to remove the expected variation due to changes in average clone size. Inset, dependence of the cut-off frequency *x*_c_ on the exponent *p* when frequency spectra are approximated by a power law *f*(*x*) *∝ x*^*p*^ for *x ≤ x*_c_ and *f*(*x*) = 0 for *x > x*_c_. Dots show the cutoff estimated numerically as the value for which the second derivative of the rescaled curves first drops below −4, plotted against the observed exponents *p* ≈ {−0.9*, −*0.72*, −*0.47, 0, 1} for the small-frequency behaviour.

To uncover variations due to long-range dispersal beyond changes in the average clone size, we rescaled the frequency spectra by the expected dependence on *X*_ave_, which we have already established as being set by the mutation-expansion balance via the characteristic size *χ*. To establish the form of this rescaling, we assume that for a given dispersal kernel, soft sweep patterns at different mutation rates are self-similar when distances are rescaled by the characteristic length *χ*. Under this assumption, the probability distribution of clone sizes in an infinitely large range is a function only of the rescaled clone mass *s* ≡ *X/X*_ave_; i.e. the probability of finding a clone between *s* and *s* + *δs* is *P*_*µ*_(*s*) *δs*, where the density function *P*_*µ*_ depends only on the dispersal kernel and not on the rescaled mutation rate.

For finite ranges of extent *L* much larger than *χ*, we can now express the average allele frequency spectrum in terms of *P*_*µ*_. The expected number of unique alleles in the range is *L*^*d*^*/X*_ave_. Within these alleles, the probability of finding an allele in the frequency range (*x, x* + *δx*) is *P*_*µ*_(*L*^*d*^*x/X*_ave_) *× L*^*d*^*δx/X*_ave_. Therefore, the expected number of alleles with frequencies between *x* and *x* + *δx is*

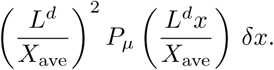

Upon comparing this expression the definition of the allele frequency spectrum for the finite range, we arrive at

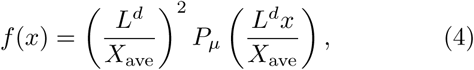

Eq. 4 implies that for a given dispersal kernel, the dependence of the allele frequency spectrum on mutation rate and range size is completely captured by the ratio *L*^*d*^*/X*_ave_. In particular, when *f*(*x*) is multiplied by (*X*_ave_*/L*^*d*^)^2^ and the frequency by *L*^*d*^*/X*_ave_, frequency spectra for different values of 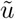 ought to collapse onto a single curve for each *µ*. Fig. 6(c) shows that upon such a rescaling (with ⟨*X*⟩ used as a simulation-derived estimate of *X*_ave_), curves for the same value of *µ* from panels (a) and (b) largely coincide, confirming that most of the dependence of the frequency spectrum on mutation rate is captured by the variation of the single length scale *χ* and, through it, the expected clone mass *X*_ave_. Note that we can use the fact that *X*_ave_ ∝ *χ*^*d*^ with a kernel-independent prefactor to rewrite Eq. 4 as *f*(*x*) = (*L/χ*)^2*d*^*G*_*µ*_(*L*^*d*^*x/χ*^*d*^), where *G*_*µ*_ is independent of 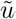, which explicitly shows the role of *χ* in scaling the allele frequency spectrum.

The arguments leading to Eq. 4 relied on the assumption that only one characteristic scale exists for the soft sweep patterns. For our class of kernels, this assumption is only exact in the regime of power-law growth, for which the halo extent scale *ψ* is proportional to *χ*. In the stretched-exponential and marginal growth cases, by contrast, *ψ* acts as an independent length scale from *χ* with its own mutation-rate dependence. In Appendix E, we show that the consequent corrections to Eq. 4 are weak (logarithmic in mutation rate and system size) and are strongest when *µ* approaches 0, validating the effectiveness of the proposed rescaling over all regimes away from the well-mixed limit.

The scaled frequency spectra show that broader dispersal kernels favour broader allele frequency spectra even after accounting for changes in the average clone size. At *µ* = 4, the steep decline in the frequency spectrum occurs near the frequency expected of an average clone, *x* ≈ 〈*X*〈/*L*. As *µ* is reduced, the falloff occurs at higher frequencies; at *µ* = 0.4, for instance, clones with frequencies an order of magnitude higher than the average clone are still likely. Qualitatively, this trend is a result of the increased nonlinearity of the growth functions *ℓ*(*t*) for broader dispersal. If we assume no interference among distinct clones until the time *t*^*\**^, the size of an allele which arrives at time *t*_*i*_ is proportional to *ℓ*^*d*^(*t*^*\**^*-t*_*i*_). For a given spread of arrival times of mutations, the spread of final clone sizes is significantly enhanced by nonlinearity in *ℓ*(*t*). Therefore, the increased departure from linear growth in *ℓ*(*t*) as *µ* → 0 gives rise to broader clone size distributions. Deterministic approximations to the clone size distributions expected for the asymptotic *l*(*t*) forms in 1D, described in Appendix E, support this heuristic picture.

Although we do not have analytical expressions for the frequency spectra at intermediate *µ*, the measured curves and deterministic calculations suggest a simple approximate form for the allele frequency spectra: extend the power-law behaviour observed at intermediate frequencies [straight parts of the curves in Fig. 6(a)–(b)] from *x* = 0 up to a cutoff frequency corresponding to the location of the sharp dropoff in *f*(*x*). Quantitatively, we consider an ansatz for the frequency spectra with two parameters:

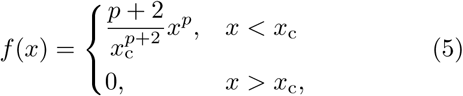

i.e. a power-law behaviour characterized by exponent *p*, up to some maximal frequency *x*_c_, with the constant of proportionality determined by the normalization. The values *p* and *x*_c_ are determined from the numerical data, but are also consistent with theoretical arguments (Appendix E). The small-*x* behaviour of the two limiting spectra, *f*_*∞*_(*x*) *∼ x*^−1^ and *f*_w_(*x*) *∼ x* as *x →* 0, imply that *p* is restricted to vary from −1 to 1 as *μ* increases from zero. Despite its simplicity, this approximation can be used to quantify the relationships among various features of the clone size distributions as we show in Appendix F. For instance, the power-law ansatz predicts a relation between the average clone size and the cut-off frequency, *Lx*_c_*/X*_ave_ = (*p* + 1)*/*(*p* + 2), which matches the trends observed in the rescaled frequency spectra, see inset to Fig. 6(c).

#### 1. Global sampling statistics

The utility of *f*(*x*) in the context of soft sweep detection is made apparent by noting that the probability *P*_hard_(*j*) of finding only one unique allele in a sample of size *j* ≥ 2 drawn randomly from the population (i.e. detecting a *hard* sweep) is [18]

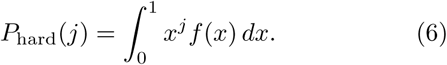

The probability of observing a *soft* sweep in a sample of size *j* is simply *P*_soft_ = 1 – *P*_hard_(*j*). (Although *P*_soft_ might be more relevant to soft sweeps, we deal with *P*_hard_(*j*) in the following sections because it is more straightforward to compute and manipulate mathematically.) Since *xf*(*x*) does not diverge as *x* → 0 for all observed frequency spectra, the integral in Eq. 6 is dominated by contributions from the high-frequency region of *f*(*x*) and is therefore highly sensitive to the breadth of the frequency spectrum. Using the power-law ansatz for the frequency spectrum, Eq. 5, gives 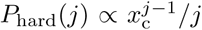 for large *j* (see Appendix F): the dominant behaviour is an exponential decay with sample size, with the decay scale set by the high-frequency cutoff *x*_c_. At a given rescaled mutation rate, this cutoff falls by many orders of magnitude as *μ* is increased, as we saw in Fig. 6(a)–(b). As a consequence, the probability of finding a monoallelic sample also falls dramatically with increasing *μ*, see Fig. 7(a). Analytical calculations of *P*_hard_ using *f*_*∞*_(*x*) and *f*_w_(*x*) in the *μ* → 0 and *μ* ≫ *d* + 1 limits (dashed and dash-dotted lines) provide bounds on the variation (see Appendix G for explicit forms of *P*_hard_(*j*) in these limits).

**FIG. 7.**
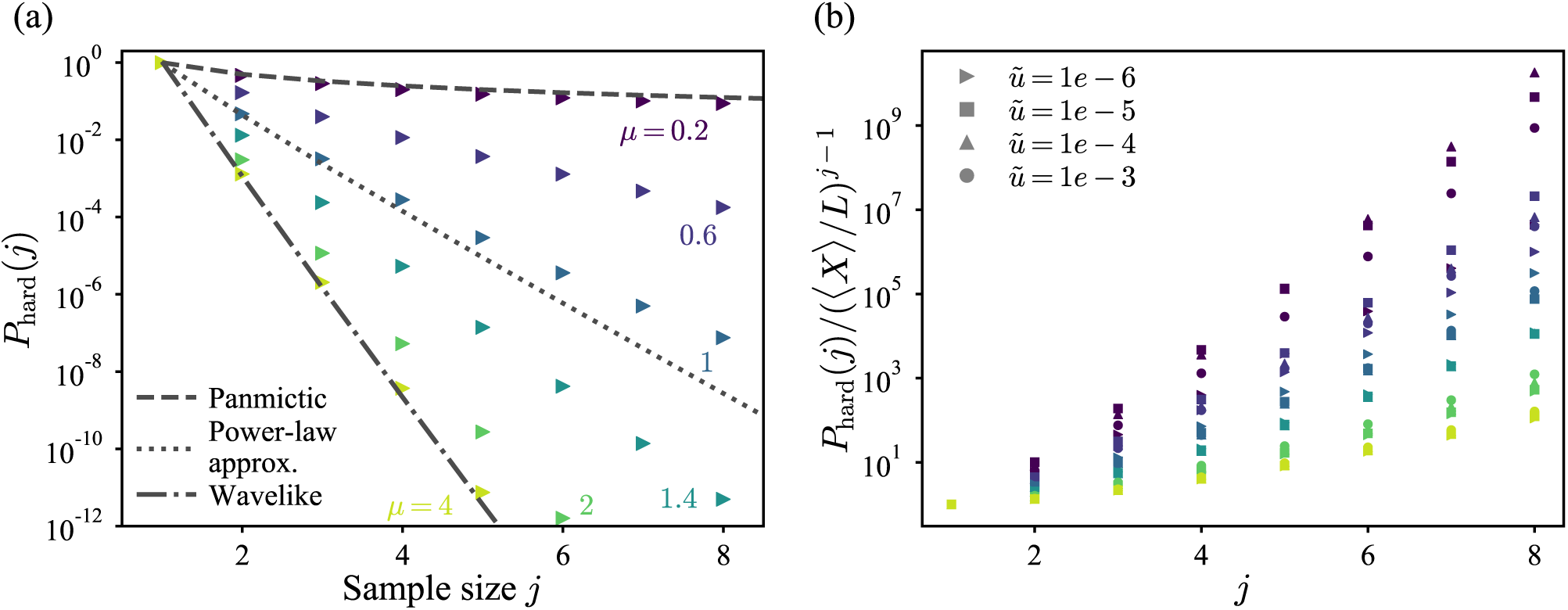
Probability of observing only one allele in a finite sample. (a) The probability *P*_hard_(*j*) of observing a hard (i.e. monoallelic) sweep in a sample of size *j* chosen at random from the range, computed from simulated clone size distributions for different dispersal kernels (colours, labeled) with *L* = 10^6^ and 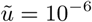 in 1D. Three analytical forms are shown as dotted lines (from top to bottom): the Ewens’ sampling result for the panmictic case, the approximate form derived using a hard-cutoff ansatz for the allele frequency spectrum for *μ* = 1 (Eq. F3), and *P*_hard_ calculated from the exact *f*(*x*) for the wavelike spreading limit (Appendix G). (b) The same quantity computed across a range of rescaled mutation rates (symbols), and scaled by the expectation for a range with all clones having the same size and hence the same frequency *X*_ave_*/L*.

We have seen that increased long-range dispersal broadens the frequency spectrum both by increasing the average clone size, and by enhancing the spread of clone sizes around the average. In the hypothetical case of all clones having the same size *X*_ave_, a monoallelic sample of size *j* is obtained by having the last *j* – 1 samples drawn from the same clone as the first sample, which occurs with probability (*X*_ave_*/L*^*d*^)^*j*–1^ ∝ (*χ/L*)^*d*(*j*–1)^. To distinguish the effect of the shape of *f*(*x*) from that of the average size of clones, we scale *P*_hard_(*j*) in 1D simulations by (⟨*X*⟩*/L*)^*j*–1^ for a range of rescaled mutation rates, see Fig. 7(b). (As before, we use ⟨*X*⟩ for a simulation-derived estimate of *X*_ave_.) If the sampling statistics were determined primarily by the average clone size (which in turn is set by *χ*) and the effect of variations in the shape of *f*(*x*) were insignificant, we would expect the rescaled *P*_hard_(*j*) for different kernels to all collapse on the same curve. Instead, we find that the sampling statistics vary significantly with *μ* even when accounting for differences in average clone size. Whereas the rescaling captures a significant amount of the variation in *P*_hard_(*j*) *within* each value of *μ* (with a residual 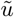-dependence that differs for the different regimes of *ℓ*(*t*), and is due to the relevance of the additional length scales outside the power-law growth region), the rescaled curves vary widely among the different dispersal kernels.

Fig. 7(b) quantifies the influence of long-range dispersal on soft sweep detection beyond merely setting the average size of clones: if mutation rates are adjusted so that the characteristic length scales *χ* and hence the average clone sizes are comparable for different dispersal kernels, soft sweeps continue to be *less* likely to be detected for broader kernels (smaller *μ*). This happens because the range has a larger contribution from high-frequency clones with *x >*(*χ/L*)^*d*^ for broader dispersal kernels, making monoallelic sampling more likely. In summary, not only does broadening the dispersal kernel make sweeps harder, it also makes their *detection* less likely. Since a wide range of possible outcomes separates the two limits of panmictic (*μ* → 0) and wavelike spreading (*μ ≫ d* + 1), predictions based on these extremes might perform poorly in making inferences from sampling statistics in populations with intermediate long-range dispersal.

### E. Local sampling protocols are highly sensitive to the core-halo structure

Population genomic studies are often limited not only in the number of independent samples available, but in their geographic distribution as well. Samples tend to be clustered in regions chosen for a variety of reasons such as anthropological or ecological significance, or practical limitations. The analysis of the last section would apply to comparing samples across different regions, provided that these are relatively well spread out in the range. Here we focus on the variation within local samples from a subrange of the entire population. As illustrated by the wide variation in local diversity within the highlighted subranges (dashed boxes) in Fig. 1(a), inferences based on local sampling can be significantly different from inferences based on global information, and may be very sensitive to modes of long-range dispersal.

Long-range dispersal enhances local diversity. When clones extend over a much wider spatial range than required by their mass (Fig. 5), local subranges contain alleles whose origins lie far away from the subrange, and are consequently more diverse than expected from the diversity of the range as a whole. To quantitatively illustrate this effect, we compute sampling statistics for different dispersal kernels and subrange sizes from 1D simulations with a global range size much larger than the characteristic length scale *χ* (Fig. 8). (Subrange size, denoted by *L*_s_, and extent are equivalent in our 1D simulations.) We observe that the smaller clones expected at higher values of *μ* favour the detection of soft sweeps globally (Fig. 8a), but the diversity is less detectable in samples from subranges that are smaller than the characteristic size shared by the compact domains at *μ* = 4. By contrast, samples from smaller subranges continue to show signatures of soft sweeps for broader dispersal kernels (Fig. 8b–c).

**FIG. 8.**
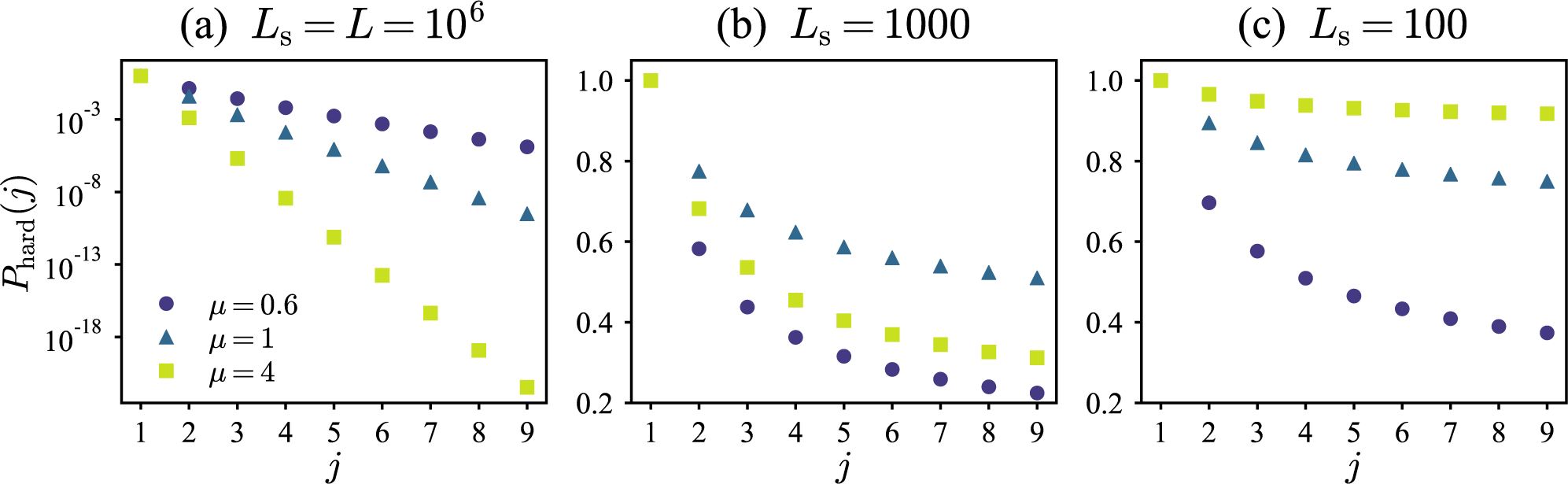
Dispersal promotes soft sweep detection in small subranges. (a)–(c) Probability of observing a hard sweep in *j* samples randomly chosen from contiguous subranges of different sizes *L*_s_ in simulated 1D ranges of size *L* = 10^6^, with rescaled mutation rate 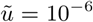. At the same mutation rate, broader dispersal kernels lead to a larger average clone size (⟨*X*⟩ 980, 1.6 *×* 10^4^, 4 *×* 10^4^ for *μ* ={4, 1, 0.6} respectively), which reduces the total number of alleles and favours hard sweep signatures when the sampling is done over the entire range [*L* = *L*_s_, (a)]. However, when *L*_s_ is reduced [(b)–(c)], the detection of soft sweeps become increasingly likely for the broader dispersal kernels; the broken-up structure of clones compensates for their smaller overall number. For small enough subranges, the order of values of *P*_hard_(*j*) with increasing *μ* is inverted compared to the values at *L*_s_ = *L*.

To compare the sensitivity of soft sweep detection to subrange size across different dispersal kernels and mutation rates, we focus on the probability of detecting the same allele in a *pair* of individuals randomly sampled from a subrange, *P*_hard,s_(2) (also called the species homoallelicity of the subrange). This probability is high only when the subrange is mostly occupied by the *core* of a single clone; it is low if the subrange contains cores belonging to different clones, or a combination of cores and haloes. Therefore, we expect *χ* (or equivalently the average mass-equivalent radius ⟨*r*_eq_⟩, which we may use as a simulation-derived estimate for *χ* in 1D) to also be the relevant scale to compare *L*_s_ values across different situations. Fig. 9(a) shows the dependence of *P*_hard,s_(2) on *L*_s_*/* ⟨*r*_eq_⟩ for different dispersal kernels and mutation rates in the *χ ≪ L* limit. As with the global sampling probabilities reported in Fig. 7(b), we find that the rescaling of subrange size with *r*_eq_ captures much of the variation among different mutation rates (symbols) for a given dispersal kernel. In contrast with the global sampling statistics, however, hard sweep detection probabilities are suppressed (or equivalently, soft sweeps are *easier* to detect for the same rescaled subrange size) as the jump kernel is broadened. At high values of *μ* in the wavelike expansion limit, the shape of the curves is well-approximated by the null expectation for an idealized clone size distribution where all clones are perfectly contiguous segments of equal size *X*_ave_. As *μ* falls below *d* + 1, the prevalence of overlapping haloes increases local diversity at the scale of satellite clusters, much smaller than the typical clone size would dictate. The effect is especially strong in the marginal and stretched-exponential growth regimes (*μ ≤ d*), which was associated with the halo dominating over the core (Figs. 3 and 5).

**FIG. 9.**
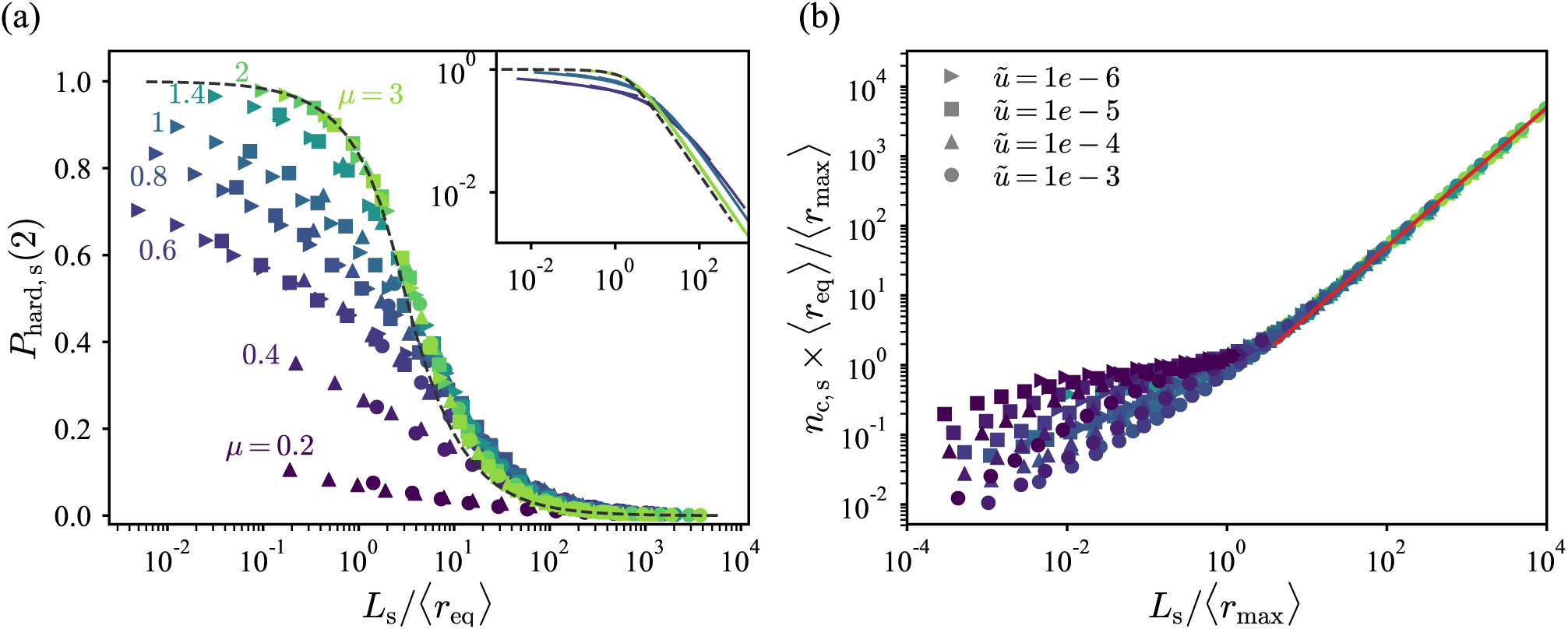
Different measures of diversity within a subrange are sensitive to different characteristic scales. (a) Probability *P*_hard,s_(2) of observing a single allele in a pair drawn from a subrange of size *L*_s_ for different dispersal kernels (colours, labeled) and mutation rates [symbols, see legend in panel (b)], for 1D simulations with *L* = 10^6^, as a function of the ratio *L*_s_*/X*_ave_. In all cases, the population range was chosen to be many times larger than the characteristic size *χ* and harbours many distinct alleles. The dashed line is the prediction if all clones are of the same size *X*_ave_, in which case geometry dictates that *P*_hard,s_(2) = {1 – *x*/3, *x* < 1; 1/*x* – 1*/*(3*x*^2^)*, x* ≥ 1}. The inset shows data for *μ* = {0.6, 1.0, 3.0} on log axes. (b) Number of distinct alleles *n*_c,s_ observed in a subrange of size *L*_s_, shown as a function of the ratio *L*_s_*/* ⟨*r*_max_⟩. Values are scaled by ⟨*r*_max_⟩*/* ⟨*r*_eq_⟩, the expected number of clones in the area occupied by the average halo. The solid line corresponds to *n*_c,s_ ⟨*r*_eq_⟩*/* ⟨*r*_max_⟩ = *L*_s_*/*(2 ⟨*r*_max_⟩), or equivalently *n*_c,s_ = *L*_s_*/*(2 ⟨*r*_eq_⟩).

A different measure of subrange diversity is the total number of distinct alleles present in a subrange on average, which we call *n*_c,s_. Unlike the subrange homoallelicity, which was dominated by the most prevalent clone in the subrange, this measure gives equal weight to all clones, and is sensitive to haloes that overlap with the subrange. The expected number of distinct cores in the subrange is *L*_s_*/*(2 ⟨*r*_eq_⟩); in the absence of haloes, we would expect *n*_c,s_ to be equal to this value. How-ever, haloes of clones whose cores are outside the sub-range would cause *n*_c,s_ to exceed the number of cores in the subrange. This enhancement in diversity due to encroaching haloes would be expected to occur only when the subrange is smaller than the average clone extent including the halo, i.e., when *L*_s_ *<* 2 ⟨*r*_max_⟩. When the subrange is larger than the typical halo extent, the cores whose haloes contribute to *n*_c,s_ are also expected to lie within the subrange, and are accounted for in *L*_s_*/*(2 ⟨*r*_eq_⟩). This expectation is confirmed in Fig. 9(b). When the subrange size is rescaled by the extent of the clone including the halo, the average number of distinct alleles in the subrange follows *n*_c,s_ = *L*_s_*/*(2 ⟨*r*_eq_⟩) (solid line) in all cases, provided *L*_s_*/* ⟨*r*_max_⟩ *>* 2. For smaller subrange sizes, *n*_c,s_ lies above this estimate, reflecting the enhancement of local diversity due to encroaching haloes.

## III. DISCUSSION

Adaptation in a spatially extended population often uses different alleles in different geographic regions, even if the selection pressure is homogeneous across the entire range. The probability of such *convergent adaptation* [21] and the patterns of spatial soft sweeps that result depend on two factors: the potential for the population to recruit adaptive variants from either new mutations or from the standing genetic variation, and the mode of dispersal. Previous work has focused on the two extremes of dispersal phenomena: panmictic populations without spatial structure [3, 4, 5] or wavelike spreading due to local diffusion of organisms [11, 21]. However, gene flow in many natural populations does not conform strictly to either limit. Many species experience some long-distance dispersal either through active transport or through passive hitchhiking on wind, water, or migrating animals including humans [12, 13, 14]. The dynamics of adaptation of populations with a large range can be strongly influenced by long-distance dispersal even when dispersal events are rare [22].

We have described spatial patterns of convergent adaptation for a general dispersal model, with jump rates taken from a kernel that falls off as a power-law with distance. Although the underlying analysis is applicable to more general dispersal kernels, our specific choice of kernel allows us to span a wide range of outcomes using a single parameter. We have shown that long-range dispersal tends to break up mutant clones into a core region dominated by the clone, surrounded by a disconnected halo of satellite clusters and isolated demes which mingle with other alleles. A key result of our analysis is that although the total mass of a clone is well-captured by the extent of the core region, the sparse halo can extend out to distances that are significantly larger than the core, sometimes by orders of magnitude. Therefore, understanding clone masses alone provides incomplete information about spatial soft sweep patterns, and can vastly underestimate the true extent of mutant clones.

By analyzing the balance between the jump-driven expansion of solitary clones and the introduction of new mutations, we have identified three characteristic length scales that quantify the spatial relationships between core and halo: the characteristic core extent *χ*, which sets the average clone mass; the radial extent *ψ* within which well-developed satellite clusters are expected; and the outer limit *ζ* within which both satellite clusters and isolated demes are typically found. As the kernel exponent *μ* is varied, these length scales demarcate three regimes with qualitatively different core-halo relationships: compact cores with insignificant haloes, similar to the case of wavelike growth, for *μ > d* + 1; a dominant high-occupancy core surrounded by a halo of well-developed satellite clusters which extend to a size-independent multiple of the core radius (*ζ ∼ ψ ∝ χ*) when *d* < *μ* < *d* + 1; and a halo including a significant number of isolated demes in addition to satellite clusters, which may extend over a region orders of magnitude larger than the core (*ζ ≫ ψ ≫ χ*) when *μ < d*.

We have also studied the signatures left behind by these patterns on population samples that are taken either from a local region, or globally from the entire range. Under which conditions, and for which types of samples, can we expect to observe a soft sweep? We have found that when ranges with similar overall diversity (as judged by the number of distinct clones in the entire range) are compared, broadening the dispersal kernel has opposing effects on soft sweep detection at global and local scales: soft sweeps become harder to detect in a global random sample, but easier to detect in samples from smaller subranges.

Besides having consequences for detecting and interpreting evidence for spatial soft sweeps, the breakup of mutant clones by long-range dispersal also impacts future evolution after the soft sweep has completed. Our analysis describes the spatial patterns arising in the regime of strong selection, where the large advantage of beneficial mutants over the wildtype dominates the evolutionary dynamics. Once the entire population has adapted to the driving selection pressure, smaller fitness differences among the distinct alleles will become significant, and modify the spatial patterns on longer time scales. Selection is most sensitive to these fitness differences at the boundaries separating demes belonging to different clones. For the same global diversity, the total length of these boundaries is strongly influenced by the connectivity of clones, and grows significantly as the kernel exponent is reduced, thereby modifying the post-sweep evolution of the population. The post-sweep evolution could also favour well-developed satellite clusters over isolated demes of one allele within a region dominated by another: isolated demes are likely to be taken over by their surrounding allele through local diffusion of individuals. Therefore, the characteristic length *ψ* may prove to be a relevant spatial scale for the post-sweep evolution, even in the regime *μ < d* where *ζ* sets the extent of the halo in the sweep patterns.

Although a quantitative evaluation of our model using real-world genomic data is beyond the scope of this work, some qualitative features of long-range dispersal can be identified in previous studies of spatial soft sweeps. The evolution of resistance to widely-adopted drugs in the malarial parasite *Plasmodium falciparium* is a well-studied example of a soft sweep arising in response to a broadly applied selective pressure. While multiple mutant haplotypes conferring resistance to pyrimethamine-based drugs have been observed across Africa and South-east Asia, the number of distinct haplotypes is smaller than would have been expected if resistance-granting mutations were confined to their area of origin [23]; this feature has been linked to long-distance migration of parasites through their human hosts, which allowed individual haplotypes to quickly spread across disconnected parts of the globe [24]. Within the same soft sweep, high levels of spatial mixing of distinct resistant lineages was also observed in some sub-regions [25]. These observations are consistent with the contrasting effects of long-range dispersal we have quantified in our model: at a given rescaled mutation rate, dispersal reduces diversity globally, but increases the mixing of alleles locally. Advances in sequencing technology have driven rapid improvements in the spatiotemporal resolution of drug-resistance evolution studies [26], making them a promising candidate for quantitative analysis of the spatial soft sweep patterns we have described.

Many interesting questions remain to be explored. Our simulation studies in *d* = 2 could be significantly expanded. We have also focused on the limit in which the average clone size is many times smaller than the entire range. It would also be interesting to study the statistics of soft sweeps when the extent of the range is comparable to the characteristic length scale *χ*, making a soft sweep an event of low but significant probability which may vary significantly with the dispersal kernel.

The applicability of our results to continuous populations without an imposed deme structure is an open problem. In our model, the deme structure is used to impose a local population density and allows us to separate the local dynamics of fixation from the large-scale behaviour driven by rare but consequential jumps. However, the theoretical picture of growth via the merger of satellite outbreaks with an expanding core does not rely on the deme structure. Therefore, we expect aspects of our results to also hold in continuous populations under certain parameter regimes. However, explicitly translating the parameters and defining the correct continuum limit of deme-based models is known to be challenging [27], and presents an interesting avenue for future work. Our simulations could also be modified to exploit advances in computational modeling of continuum populations [28].

The model can also be extended to include additional mechanisms involved in parallel adaptation. Besides recurring mutations, standing genetic variation (SGV) in the population is a important source of diversity for soft sweeps [3]. Long-range dispersal could impact both the spatial distribution of SGV before selection begins to act, and the spreading of alleles from distinct variational origins during the sweep [21]; both situations can be explored through extensions of our model. In the latter case, we expect the distinct regimes of core-halo patterns for different jump kernels to persist, but with the characteristic core size set by the initial distribution of variational origins rather than mutation-expansion balance.

The necessity of including heterogeneity motivates a natural set of extensions of the model. When soft sweeps arise due to mutations at different loci producing similar phenotypic effects, some variation in fitness among the distinct variants is inevitable. In panmictic models, fitness variations do not significantly affect the probability of observing a soft sweep, provided that the variations are small relative to the absolute fitness advantage of mutants over the wildtype [5]. Since spatial structure restricts competition to the geographic neighbourhood of a clone, we expect the effect of fitness variation to be even weaker than for panmictic populations, and our results should be robust to a small amount of variation in fitness effects. However, when fitness variations among mutations are large enough to be significant, the impact of the variations could depend on the dispersal kernel, and show qualitatively different behaviours in the distinct regimes of power-law and stretched-exponential growth. Similarly, spatial heterogeneities in the selection pressures could lead to so-called “patchy” landscapes which lead to certain mutations being highly beneficial in some patches but neutral or even deleterious in others [29]. Convergent adaptation on patchy landscapes is likely to be significantly impacted by long-range dispersal which would allow mutations to spread efficiently to geographically separated patches.

Finally, the assumptions of strong selection and weak mutation/migration allowed us to ignore the dynamics of introduction of beneficial mutations within a deme. Relaxing these assumptions would lead us to a more general model with an additional time scale characterizing the local well-mixed dynamics at the deme level. The interplay between this time scale and the time scales governing the large-scale dynamics driven by long-range dispersal could lead to new patterns of genetic variation during convergent adaptation.

## IV. MATERIALS & METHODS

### A. Simulation methods

Simulations were written in the C++ programming language, and utilized the standard Mersenne Twister engine to generate pseudorandom numbers. A simulation of linear size *L* in *d* dimensions is begun by initializing an array of integers of size *L*^*d*^. Each array position corresponds to a single deme, and the associated integer value stores the allelic type. The array is initialized with all demes bearing the value 0 signifying the wildtype (WT).

As described in the text, the simulations only need to incorporate the two types of events which could potentially change the identity of a deme: a mutation of a WT deme, or an attempted migration from a mutant deme. To accomplish this, each deme is assigned a weight of 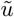 if WT, and 1 if a mutant deme. At each discrete simulation step, a deme is picked at random with probability proportional to its weight. If the deme chosen is WT, it is assigned a unique integer that was not previously present in the array. If the deme chosen contains a mutant allele, a jump is attempted. The jump distance *r* is obtained by drawing a random number *X* evenly distributed between 0 and 1, and computing the variable *r* = *X*^−1*/µ*^; this produces a variable with normalized probability density function *P*(*r*) = *µr*^−(1+*µ*)^ for kernel exponent *µ*. The distance is then multiplied with a random *d*-dimensional unit vector (simply *±*1 in *d* = 1, and evenly distributed on the unit circle in *d* = 2). Each vector component is rounded to the nearest integer to obtain a jump vector on the lattice. The target position for the migration attempt is obtained by adding this jump vector to the source position, and wrapping the result into the range of size *L*^*d*^ assuming periodic boundary conditions.

If the target deme is WT, its value is updated with the allelic identity of the source; otherwise the migration attempt is unsuccessful. If the simulation step ends in a mutation or a successful migration, the probability weights associated with the demes are updated and the next step is executed. The simulation continues until all *L*^*d*^ array positions contain nonzero integers signifying the completion of the sweep. The final array of *L*^*d*^ integers constitutes the simulation output.

A single simulation took between a few minutes and 24h of CPU time depending on the parameter values. Simulation results were processed using scripts written in the Python programming language. All reported results were obtained by averaging over 20-100 independent simulations for each set of parameters, depending on system size.

## ACKNOWLEDGMENTS

The authors thank Graham Coop and the reviewers for valuable feedback during the review process. JP thanks Diana Fusco and Benjamin H. Good for insightful discussions. Research reported in this publication was supported by a National Science Foundation Career Award (#1555330) and by a Simons Investigator award from the Simons Foundation (#327934). This research used the Savio computational cluster resource provided by the Berkeley Research Computing program at the University of California, Berkeley (supported by the UC Berkeley Chancellor, Vice Chancellor for Research, and Chief Information Officer); and resources of the National Energy Research Scientific Computing Center (NERSC), a U.S. Department of Energy Office of Science User Facility operated under Contract No. DE-AC02-05CH11231.

## Appendix A: Forms for *ℓ*(*t*) and *χ*

Here we describe the analytical forms for *ℓ*(*t*) used to compute the predictions for the characteristic length scale *χ* in main text Fig. 4. Ref. 17 derived asymptotic growth forms for the long-time limit of the domain core *ℓ*(*t*) (i.e. the region within which the occupancy of the range by an isolated domain is of order 1) for dispersal kernels with tails that fall off as *r*^−(*µ*+*d*)^:

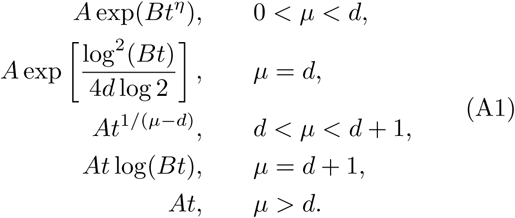

Here, *η* = log_2_[2*d/*(*d* + *µ*)], and *A* and *B* are magnitude scales for *ℓ* and *t* that depend on *µ* and on details of the dispersal kernel. (In the wavelike growth regime, *µ > d*, *A* is the front velocity of the growing domain.) The logarithmic correction to linear growth for *µ* = *d* + 1 is a conjecture for *d* = 2, which is supported by simulation data.

To extract *A* and *B* for the specific kernels used here, we performed separate simulations in which domains were grown from a single seed at the origin at *t* = 0. The domains were grown up to final masses of order 10^8^ for *µ* ≤1 and 10^5^ for *µ >* 1 in 1D, and of order 10^7^ in 2D, with the background mutation rate turned off. For each value of *µ*, 20 independent simulations were performed and the mass evolution over time, averaged over the independent runs, was equated to *ω*_*d*_*ℓ*^*d*^(*t*) following our definition of *ℓ*(*t*) in the main text. The *ℓ*(*t*) thus extracted was fit to the growth forms to obtain *A* and *B*. (Given that the growth of *ℓ* with *t* can be extremely fast for *µ < d* + 1, in practice we fit the functional dependence of log *ℓ*(*t*) against log *t*, with log *A* and log *B* as free parameters.) Using the total mass as a proxy for *ℓ*(*t*) leads to an overestimate of the true size of the core, because it also counts individuals in the inevitable “halo” that exists due to jumps from the core to regions outside it during the stochastic growth process. The halo contains a fraction of the individuals in the core, which falls as *µ* increases. This correction is expected to provide a multiplicative constant of order 1 to *ℓ*(*t*), which is inconsequential to the prediction of *X*_ave_ which itself equals *χ* only up to an overall constant for each *µ*.

The asymptotic forms only agree with the measured single-allele growth profiles when *ℓ*(*t*) has grown beyond a certain size. However, this threshold size becomes extremely large (i.e. order of the simulation range or larger) for values of *µ* close to *d* [17], making the asymptotic forms of limited utility to predict *χ*. Ref. 17 also derives an analytical scaling form for the behaviour of log_2_ *ℓ*(*t*) over a much broader range of times for *µ* close to *d*, which reads

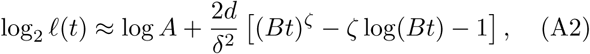

where *δ* = *µ – d* and

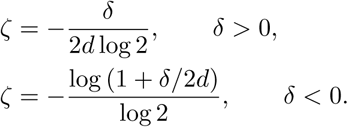

**FIG. A1.**
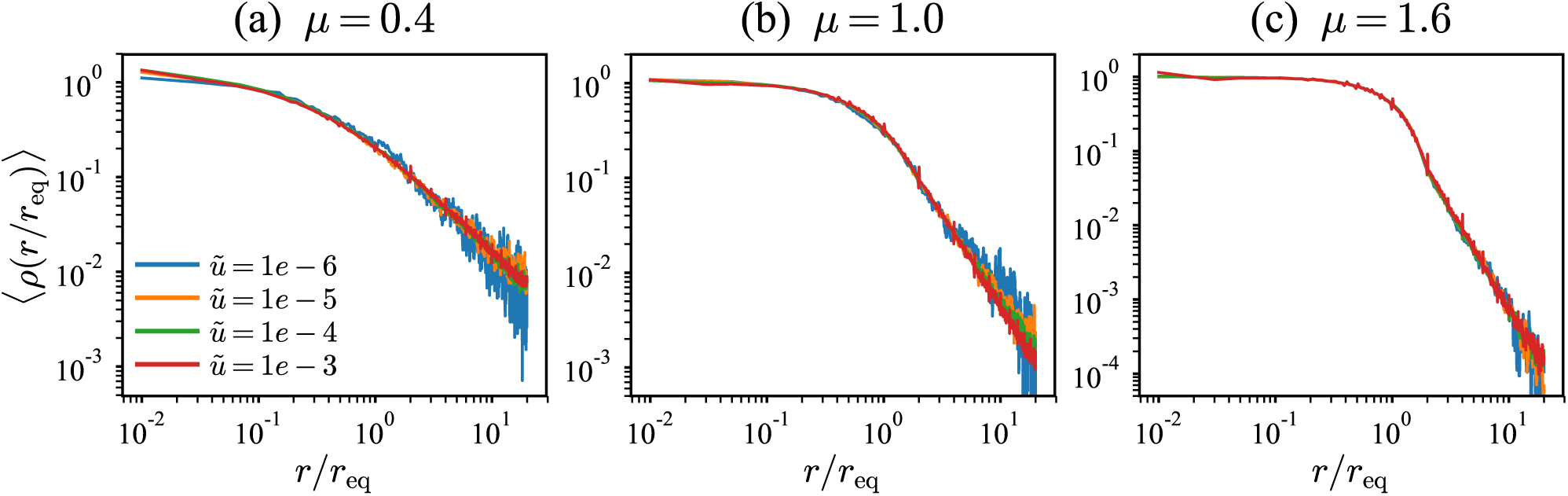
Occupancy profiles for different mutation rates collapse when the radial coordinate is rescaled by clone size. Averaged occupancy profiles ⟨*ρ*⟩(*r/r*_eq_) measured from the final states of 1D simulations with *L* = 10^6^. Panels correspond to different dispersal kernels quantified by *µ* = 0.4 (a), *µ* = 1 (b), and *µ* = 1.6 (c). Colors indicate different rescaled mutation rates. Each curve is itself an average over clones of different sizes, and the average clone sizes vary by orders of magnitude among the different values of 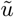. Despite this variation, the profiles for a given dispersal kernel collapse onto a single curve, confirming the validity of the rescaling of the distance variable *r* with the mass-equivalent clone radius *r*_eq_. The smallest and largest average clone sizes (at 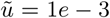 and 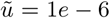 respectively) are (130, 5.8 *×*10^4^) for *µ* = 0.4; (84, 1.6 *×*10^4^) for *µ* = 1.0; and (56, 4100) for *µ* = 1.6.

As before, we used fits of log *ℓ*(*t*) against log *t* to obtain the parameter values log *A* and log *B*. From our fits to the single-allele growth simulations, we found that the scaling form is significantly more accurate than the asymptotic forms of Eq. A1 for *µ≤*1.4 in 1D, and *µ≤* 2.6 in 2D (except fo the marginal value *µ* = *d* in each case). As a result, we use the scaling form for our predictions of *χ* for these values of *µ*. Table III summarizes the values of log *A* and log *B* extracted from fits to the theoretical forms in Eqs. A1 and A2 as appropriate.

**TABLE III.**
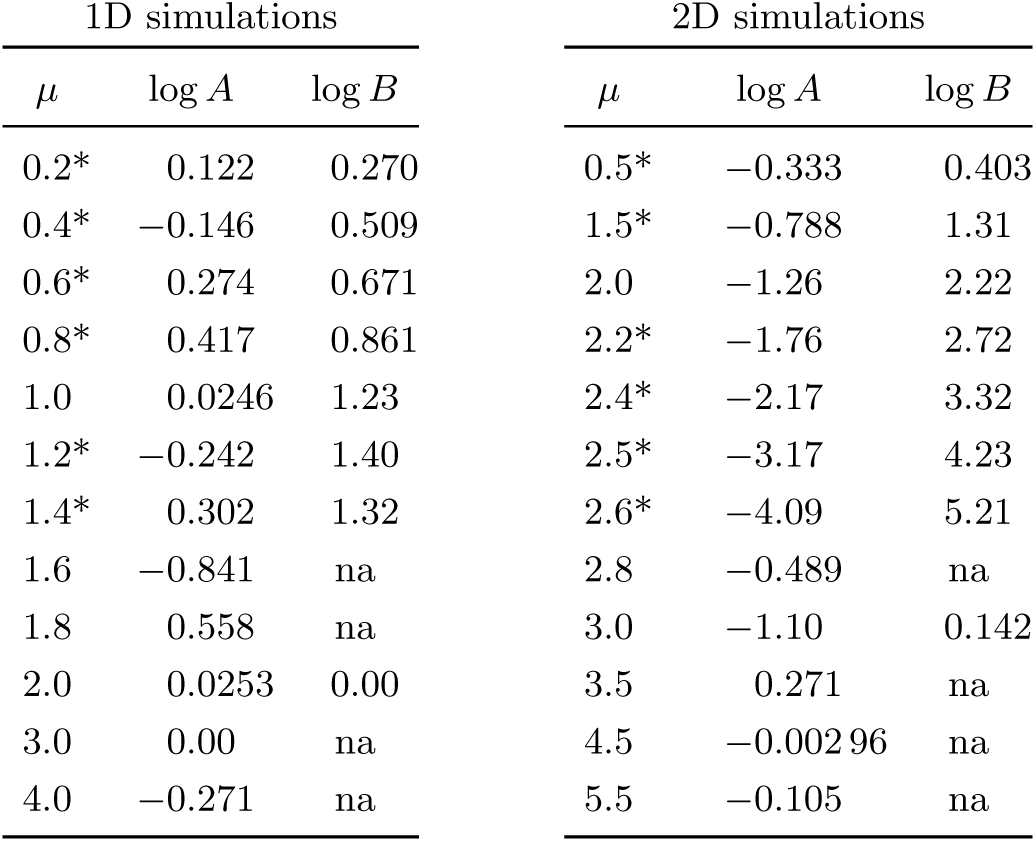
Values of parameters *A* and *B* from fits. Estimates of log *A* and log *B* obtained by fitting the growth dynamics of single clones as described in the text. The asterisk denotes use of the scaling form (Eq. A2) over the asymptotic form (Eq. A1).

In all cases, the forms for log *ℓ*(*t*) with fitted values for *A* and *B* are accurate to within a few percent for *ℓ*(*t*) of order 20 and larger. The inaccuracy of *ℓ*(*t*) for smaller domains leads to discrepancies between the measured average clone size and the prediction based on *χ*^*d*^ for large *µ* and high rescaled mutation rates, which drive down the average clone extent into the regime of inaccurate *ℓ*(*t*).

Once *A* and *B* are determined from the fit either to Eq. A1 or Eq. A2, the relation defining the characteristic length, Eq. 1 (main text), is solved to obtain *t\**(*u*) and *χ*_*µ*_(*u*) = *ℓ*_*µ*_(*t\**). Table 1 in the main text reports the functional forms for *χ* derived upon assuming that *ℓ*(*t*) follows the asymptotic forms. When the more complex scaling form is used for *ℓ*(*t*), Eq. 1 in the main text can still be solved to obtain an analytical solution for *χ*(*u*) in terms of Lambert *W*-functions. For each dispersal kernel, the solution *χ*_*µ*_(*u*) is analytically determined taking only *µ*, and the values of *A* and *B* estimated from fits (as reported in Table III) as inputs.

The characteristic length scale *χ* quantifies the balance between domain growth and mutations that sets the average domain size via *X*_ave_ ∝ *χ*^*d*^ up to a multiplicative constant of order 1; the precise relationship between *χ*^*d*^ and *X*_ave_ is determined by the distribution of domain sizes about the characteristic size, which is in turn established by the complete growth dynamics. We have an explicit form for the domain size distribution in the constant-velocity wavelike growth regime in 1D, *µ >* 2 (Eqs. D2 and D3), which allows us to derive 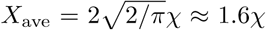 in this regime. For the 1D results in Fig. 4, we find that multiplicative constants close to 1.6 also lead to agreement between *X*_ave_(*u*) and *χ*(*u*) for other values of *µ*, over many orders of magnitude of *u*. The agreement is weakest for high *u* which corresponds to small domains (average clone sizes of 100 or smaller); here the functional forms of *ℓ*(*t*) are least accurate and stochastic effects begin to dominate the deterministic growth implied by *ℓ*(*t*).

## Appendix B: Simulation results in 2D

Here, we describe preliminary results for average clone mass, clone extent, and frequency spectra as measured from 2D simulations. Simulating large ranges is a challenge in two dimensions: effectively simulating a system in which key jumps are of order *l* in length requires a range with over *l*^2^ demes (in contrast to *l* demes in 1D). We have succeeded in simulating ranges of linear size *L* = 4096 (hence 4096^2^ ≈1.6 *×*10^7^ demes), and restricted ourselves to a range of mutation rates for which the total range mass is many times the average clone mass, so that we are in the regime of multiple-origin sweeps. However, we still expect finite-size effects to be significant for measures that depend on the spatial extent of the halo, which can stretch out to many times the mass-equivalent radius for small *µ*.

Fig. A2 compares the average clone size to the theoretical expectation *πχ*^2^, where the functions 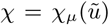 are described in Appendix A. As with the 1D results, we find quantitative agreement with the theory lines upon using a single additional parameter — an overall magnitude scale which varies between 0.75 and 0.8.

**FIG. A2.**
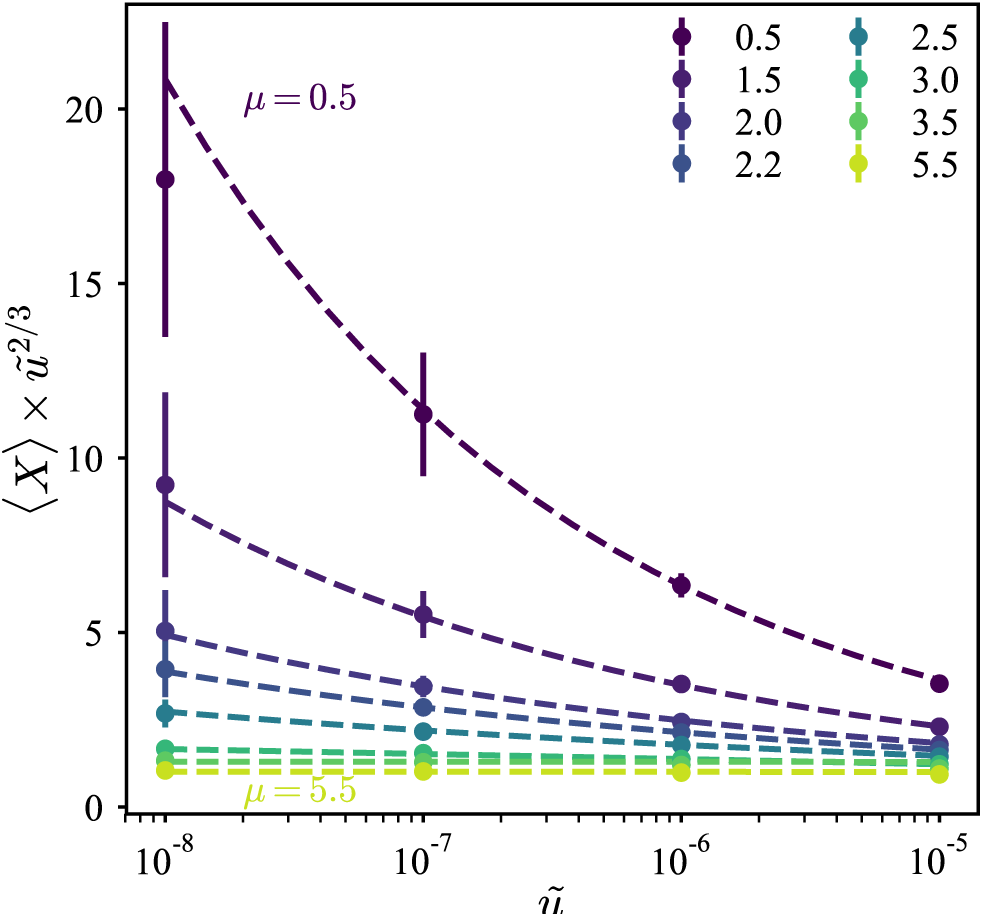
Average clone mass and mutation-expansion balance in 2D simulations. The average clone mass measured from 2D simulations as a function of rescaled mutation rate, scaled by the expected dependence 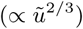 for wavelike growth. Each point represents an average over 48 independent simulations and error bars denote measured standard deviations across repetitions. Dashed lines show the theoretical prediction *πχ*^2^, using 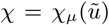 functions described in Appendix A. Each theory line is multiplied by a *µ*-dependent magnitude factor whose value is 0.8 for *µ <* 3, 0.75 for *µ* = 3, and 0.73 for *µ >* 3.

Fig. A3 reports the spatial extent of the clones from the two largest mutation rates, for which finite size effects are smallest. In 2D, we define the extent in terms of the eighth central moment: 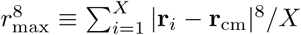 where *i* indexes the demes belonging to that clone, **r**_*i*_ is the position vector of deme *i* (computed modulo *L/*2 for each component to account for periodic boundary conditions),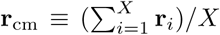 is the clone center of mass. The use of a high moment in the definition of *r*_*max*_ ensures that the farthest demes from the centre of mass contribute strongly to *r*_max_ even if they are rare. The specific choice of the eighth moment balances the need to emphasize the farthest demes (which favours a high moment) with the necessity of preventing loss of floating-point precision in the computation (which requires that the moment not be too high). Using the sixth moment leads to similar results. By contrast, using too low a moment (such as the second moment, which provides the radius of gyration of the clone) gives values of *r*_max_ that are very close to *r*_eq_ since the core provides the major contribution.

**FIG. A3.**
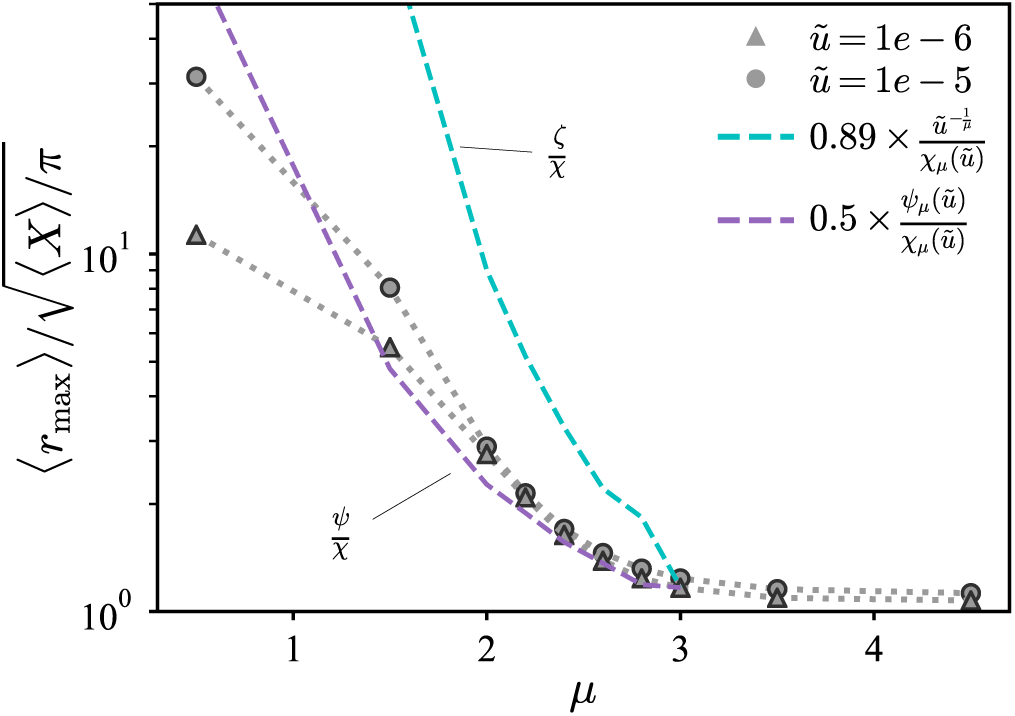
Spatial extent of clones in 2D simulations. Ensemble-averaged spatial extent of clones in 2D, normalized by the ensemble-averaged mass-equivalent radius. See text for definition of *r*_max_ in 2D. Dashed lines show theoretical expectations *ζ/χ* and *Ψ/χ* for 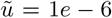, computed as described in Appendix A. The prefactor was chosen so that the lines coincide with the simulation data point at *µ* = 3. Finite size effects are more severe in 2D, and the measured values for *µ <* 2 underestimate the true values that would be measured in an infinitely large range.

We find that the dependence of the ensemble-averaged extent on the dispersal kernel is well captured by the length scale *Ψ* in the regime of power-law growth in 2D, 2 *< µ <* 3, with a single additional parameter setting the overall magnitude scale. We note that the asymptotic ratio *Ψ*_as_*/χ*_as_, which was successful in reproducing *r*_max_ for the 1D data, does *not* agree with the 2D simulation data for the current parameter range. This is because the typical sizes of clones in the 2D simulations is too small for the asymptotic growth rule (*ℓ*(*t*) ∼ *t*^1/(*µ–d*)^) to be accurate. Instead, the scaling solution from Appendix A, which accurately captures the growth of single clones at the relevant size scales, must be used.

As was seen with the 1D data, the extent starts to depart from *Ψ* as *µ*→*d*, consistent with an increased prominence of rare jumps out of the core region that land beyond well-established satellite clusters. However, the measured extent remains far below the theoretical bound *ζ/χ*, which grows extremely fast as *µ* falls below 2. We hypothesize that the ensemble averages are severely limited by finite-size effects; to attempt to match the theoretical expectation for *µ* = 1, for instance, we would require range sizes over an order of magniture larger in linear size, beyond our current capabilities for 2D simulations. Nevertheless, our limited simulations confirm that clones can attain a spatial extent many times larger than their mass-equivalent radius as the dispersal kernel is broadened.

To summarize, the results from preliminary 2D simulations show quantitative evidence for the relevance of the length scale *χ*, when combined with theoretical predictions for *ℓ*(*t*) from Ref. 17. The simulations also show that the halo can extend over much longer distances than expected for compact clone, with evidence for the relevance of the length scale *Ψ* obtained from the core-halo picture in the power-law growth regime *d < µ < d* + 1. However, more extensive simulations with much larger range sizes are needed to quantitatively test the relevance of the second scale *ζ*.

## Appendix C: Alternative derivation of secondary length scale *Ψ*

Here, we provide an alternative estimate for the length scale *Ψ* that sets the extent of the halo of a “typical” clone, which agrees with the estimate *Ψ* = *ℓ*(2*t\**) proposed in the main text. The iterative scaling picture of Ref. 17 argues that, for growth in the marginal regime near *µ* = *d*, key jumps that land at a distance *ℓ*(*t*) from the mutational origin typically occurred around time *t*/2 and spanned a distance of roughly *ℓ*(*t*) connecting source and target regions each of size ∼*ℓ*^*d*^(*t*/2) (Fig. 2b). The core extent at a given time constrains the expected number of these key jumps that have contributed to the core boundary by that time: they can be neither too rare (in which case the core would not have reached the purported boundary) nor too common (which would imply that the region should have been filled much earlier). Since the number of key jumps is itself set by the extent of the core (the source for the jumps) together with the jump kernel, the above constraint equates to a self-consistency requirement on *ℓ*(*t*) [17]:

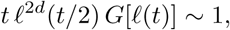

where *G*(*r*) = *J*(*r*)*r*^1*–d*^*/ω*_*d*_ is the rate of jumps per unit area of source and target regions when both are separated by a distance *r*. In the soft-sweep model, key jumps compete with new mutations in the target region, which occur at a rate of order 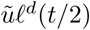. The growth of the halo is obstructed by new clones when the rate of mutations arising in the target region becomes comparable to the rate of key jumps into it from the expanding core. This requires

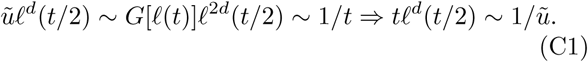

Up to factors of order unity, the above scaling relation is satisfied by *t* = 2*t\**, where *t\** was the solution to Eq. 1. Therefore, we arrive at the same expression, *Ψ=ℓ*(2*t\**), for the characteristic halo extent as we had derived in the main text from considerations of the jump-driven growth of *unobstructed* clones.

## Appendix D: Exact allele frequency spectra in the panmictic and 1D wavelike spreading limits

### 1. Panmictic limit

The panmictic limit in our lattice model would correspond to jumps being attempted from the source deme to a randomly chosen deme in the entire range. The allele frequency spectrum and related sampling probabilities can be computed exactly in this limit by mapping to an urn process. To see this, consider the evolution of allele frequencies in our lattice model when the fraction of wildtype sites is *w* and mutants occupy the lattice with individual fraction *f*_*i*_ for mutant *i*. At the next time step, the probability weight associated with picking a wildtype site to introduce a new mutation is 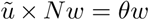, where 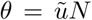 is the initial mutation rate for the empty lattice. By contrast, the probability weight associated with picking a site of mutant type *i* for an attempted dispersal event is *Nf*_*i*_, but only a fraction *w* of these attempted dispersal events is successful since the mutant only fixes in the target deme if it contains the wildtype. Therefore the probability weight of a successful reproduction of mutant *i* is *Nwf*_*i*_. The final statistics of clone sizes is determined by the *relative* rate of mutation to reproduction at each time step [5] (unlike the times for the appearance of new clones which depends on the absolute rates), which is *θ* versus *n*_*i*_ = *Nf*_*i*_ at all times since the wildtype fraction drops out.

The genealogy of new mutants in this model is identical to that of a stochastic process called Hoppe’s urn [20], which begins with an urn containing a single black ball with an assigned probability weight *θ*. At any time step, a ball is picked from the urn with probability proportional to its weight. If the black ball is chosen, it is returned along with a ball with a new colour and proportional to its weight. If the black ball is chosen, it is returned along with a ball with a new colour and probability weight 1 (a new mutant). If a coloured ball is chosen, it is returned along with one copy of itself. The relative rate of mutation to the duplication of a ball with colour *i* is *θ* versus n*_i_* at each turn, thus establishing the equivalence to our lattice model. The distributions of mutant frequencies in this urn model are the same as those for the infinite allele model at equilibrium [18]. In particular, the allele frequency spectrum is

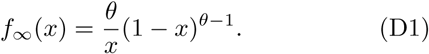

Fig. 6 shows that panmictic simulations reproduce the theoretical limit, which also persists for *µ* ≈0.5 in two dimensions.

The average clone size in the panmictic limit can be obtained from the allele frequency spectrum by computing the expected number of distinct clones *n*_*c*_. The smallest possible clone frequency is 1*/N*. Therefore, the expected number of distinct clones, *n*_*c*_, is the sum of all allowed allele frequencies, i.e. 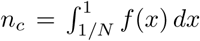, which can be evaluated exactly using *f*(*x*) from Eq. D1. For large *N*, we have 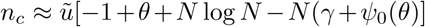, where *γ* is the Euler-Mascheroni constant and *Ψ*_0_ is the digamma function. A further simplification, valid for *θ* ≫ 1, is 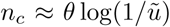 [30]. Once *n*_*c*_ is computed, the average clone size is *N/n*_*c*_.

Note that a mapping of the parallel adaptation model to an urn process was also identified in preprint [31].

### 2. Wavelike spreading limit in 1D

For *μ > d*+1, domains are predicted to grow in radially expanding waves, whose speed depends on the details of the dispersal kernel. The statistics of soft sweeps in this limit was previously explored by Ralph and Coop [11], who observed the equivalence of the process in the wavelike limit to Kolmogorov-Johnson-Mehl-Avrami (KJMA) models of grain growth. KJMA models track the evolution of isotropic domains which nucleate at random positions in space at a constant rate. Nucleated domains grow isotropically at a constant front velocity until they run into other domains, leaving a boundary separating domains that nucleated at different origins. The final pattern of domains matches the spatial pattern of clones in the mutation-expansion model, where individual domains correspond to distinct mutants.

In one dimension, the final grain size distribution for a KJMA process in which each nucleation gives rise to a unique domain is known exactly [19]. Using this result, we obtain the allele frequency spectrum for wavelike growth in 1D (*μ >* 2) as

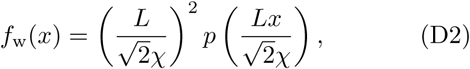

where 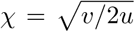 is the characteristic length scale for domains growing with front speed *v*, and

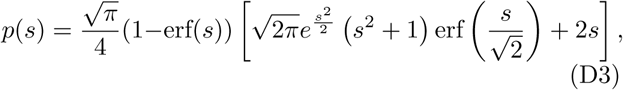

where erf is the error function. The result is valid as long as the domain sizes are not limited by the range size, i.e. *L* ≫ *χ*.

The front velocity for arbitrary *μ > d* + 1 is not known analytically, but its limiting value for very large *μ* in the lattice model is known. In the limit *μ* ≫ *d* + 1, practically all attempted jumps land exactly one lattice site away from the source (this is the lower cutoff for allowed jump distances). Isolated domains grow *via* jumps from the demes situated at the edges, only half of which are fore, the front velocity is 1/2 a lattice site per generation in the large-*μ* limit. The frequency spectra for *μ > d* + 1 approach this limit as *μ* increases, see Fig. 6. We can also extract the *μ*-dependent front speed by a one-parameter fit of Eq. D2 to the observed frequency spectra, and obtain consistent results when performing fits at different values of the mutation rate for any given *μ*, as shown in Fig. A4.

**FIG. AS4.**
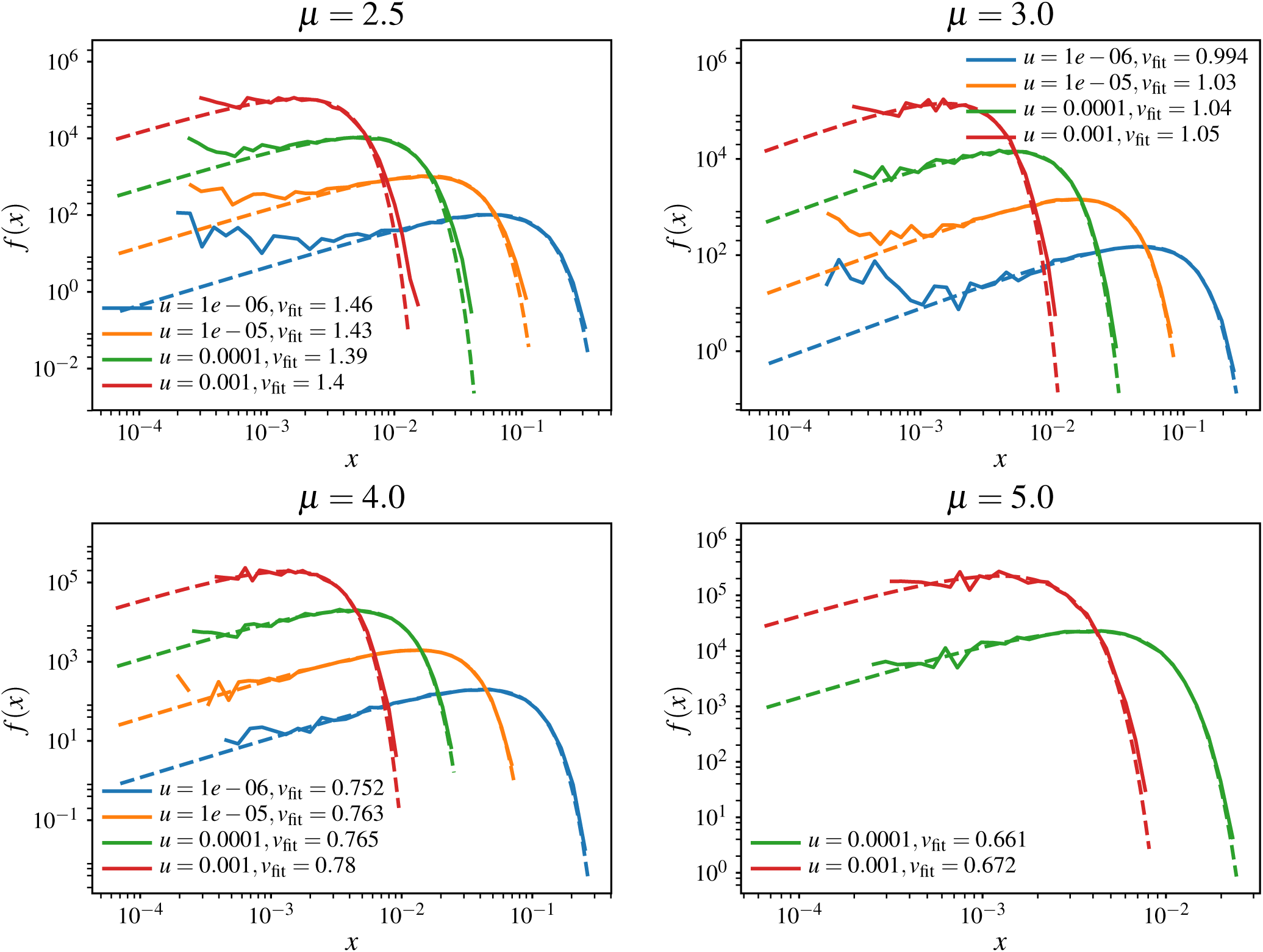
Fits to the exact frequency spectrum in the wavelike growth limit. The measured allele frequency spectra from 1D simulations in the wavelike growth regime (*µ* > *d* +1) are shown along with the theoretical form from Eqs. D2–D3. The unknown front speed *v* is extracted using a one-parameter nonlinear fit, and reported in units of lattice steps per generation. The fit values are consistent with *v* being determined by *µ* and independent of 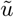. The front speed approaches the limit of 1/2 lattice steps per generation as *µ* increases.

## Appendix E: Deterministic approximation to allele frequency spectra in 1D

The analysis of the panmictic limit in the main text revealed that the distribution of alleles as *µ* → 0 was identical to that of Hoppe’s urn process. The continuous-time analogue of Hoppe’s urn is the Yule process with immigration, in which new alleles enter the population as a Poisson process with rate *θ*, and already-present individuals give birth to offspring at rate 1 without death. Yule’s process generates the same distribution of allele sizes as Hoppe’s urn, but the continuous-time description has the advantage that the dynamics of different alleles are independent: the population of allele *i* at time *t* is proportional to *e*^*t–t_i_*^ where *t*_*i*_ was the time at which it entered the population. Statistical properties of the allele frequencies, such as the frequency spectrum *f*_∞_(*x*), can be derived efficiently within this viewpoint.

In our simulations, the growth rate of alleles is *not* constant over time even if we assume panmictic migration; the success of each birth event is proportional to the wildtype fraction *w* which falls as the simulation progresses. However, as we saw in the main text, the mapping to Hoppe’s urn/Yule process remains exact because the rate of generation of new alleles is also proportional to *w* and the relative rates of birth and migration remain constant throughout the duration of the simulation in the panmictic limit. This is no longer true for *µ* > 0 when domains grow somewhat contiguously, because the likely targets for migrants become correlated with the occupancy of the lattice and the reduction in growth rate may not simply be given by the fraction *w*. If we ignore these correlations, we arrive at the following approximate continuous-time model for the establishment and growth of mutant clones: new alleles enter the population at a constant rate *θ*, and grow according to the growth rule *ℓ*(*t*) for the particular dispersal kernel, without interference from other clones.

We can make analytical headway if we further assume that the arrival of new alleles is deterministic rather than Poisson: the *k*th allele enters the population at time *t*_*k*_ = *k*/*θ*, and hence the size of the *k*th clone is *n*_*k*_ = *f*(*t*–*k*/*θ*). The total number of alleles, *K*, is fixed by the range size: 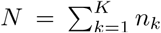. In this deterministic model, the strict time ordering of alleles implies that there are *k* alleles with size greater than or equal to *n*_*k*_; i.e. if we can invert the *n*_*k*_ relation to get *k*(*n*_*k*_), this is the survival function associated with the probability distribution of *n*_*k*_ and hence *x* = *n*_*k*_/*N*. The probability distribution of *x* is precisely the allele frequency spectrum up to a normalization.

Below, we summarize the outcome of computing *f*(*x*) according to this deterministic approximation upon using the asymptotic functional forms for *l*(*t*) in the different regimes in 1D, summarized in Table I.

### 1. Power-law growth

The deterministic approach can be used to compute an approximate frequency spectrum for the growth form *l*(*t*) = *At*^1/(*µ–*1)^, which is the asymptotic growth rule for 1 < *µ* < 2. In this case, we have a frequency spectrum that decays as a power law: *f*(*x*) ∼ *x*^*µ–*2^, up to a hard cutoff at a maximal value determined by the value of *K* that fills the entire range. Furthermore, the form admits a rescaling that ought to collapse frequency spectra across different system sizes and mutation rates: *f*(*x*) = (*L*/*X*_ave_)^2^*F*(*L*_*x*_/*X*_ave_), where *F*(*y*) = *y*^*µ–*2^ up to the cutoff *y*_max_ = *µ*/(*µ*–1), which is the same as Eq. 4 in the main text. Fig. A5 shows that the collapse works very well across different mutation rates and two system sizes. The predicted power law for *f*(*x*) is near-quantitative for all *µ* except *µ* = 1.2, which is too close to the marginal case *µ* = 1 for the asymptotic growth rule to be relevant. The predicted cutoff frequency captures the rough location of the dropoff in *f*(*x*), but the deterministic approximation fails to capture the “soft shoulder” or the clones at very large frequency, which may have an outsize influence on sampling statistics.

**FIG. A5.**
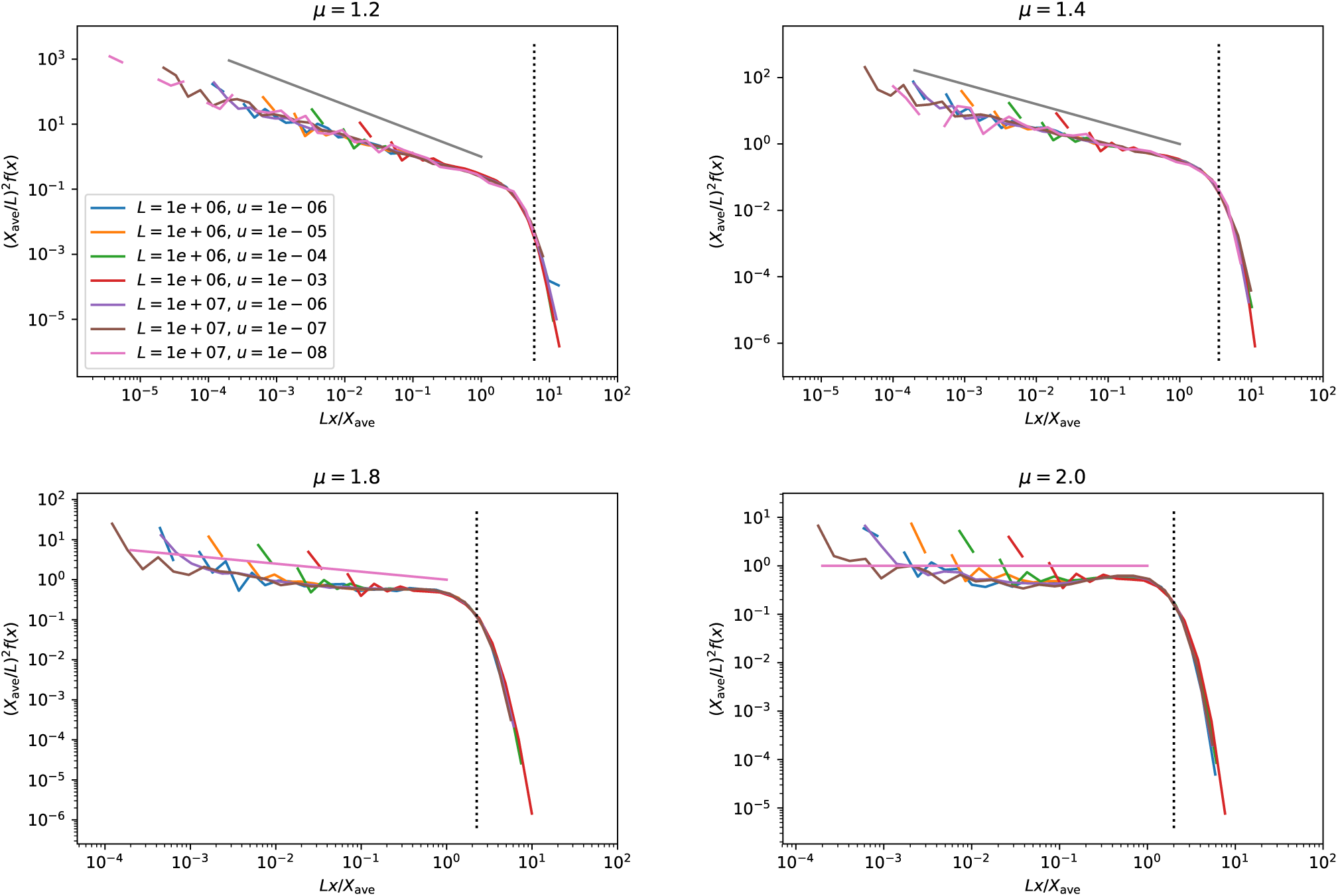
Deterministic approximation to allele frequency spectra. Allele frequency spectra in the power-law growth regime for different mutation rates and system sizes. The rescaling is suggested by the deterministic calculation, it corresponds to a clone size distribution whose only scale is the characteristic length scale *χ* or equivalently the average clone size *X*_ave_. The solid line is the prediction *f*(*y*) = *y*^*µ–*2^ and the vertical dashed line indicates the maximal rescaled allele frequency *µ*/(*µ* – 1) from the deterministic approximation.

Note that the deterministic approximation predicts a flat frequency spectrum *f*(*x*) = const. for linear growth *ℓ*(*t*) = *vt*, whereas the exact result for wavelike growth in 1D from the Axe and Yamada results, which we have seen to be quantitatively accurate for *µ* ≫ 2, predict a linear increase in the power spectrum *f*(*x*) ∝ *x* for small *x*. The difference is due to the fact that the deterministic approximation assumes that growth happens symmetrically toward both the left and the right at all times, whereas the wavelike growth limit is characterized by the left and right edges of the domain being interrupted independently as they run into other domains, so that one edge always advances for longer than the other. We can also explicitly include the log *t* correction to linear growth exactly at *µ* = 2, and we find that the low-*x* behaviour is unaffected (i.e. *f*(*x*) ∼ const. as *x* → 0) but there are contributions at higher *x*. These arise in the “shoulder” region of the spectrum, which is not captured by the deterministic analysis.

### 2. Marginal growth

If we use the growth form for *µ* = 1 in the deterministic calculation, we no longer get a simple power law for *f*(*x*); the functional form is instead 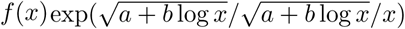 where *a* and *b* depend on the prefactors associated with *ℓ*(*t*) and on *θ* and *K*. This form is not a strict power law in *x*. However, when the various coefficients are computed using the full expression for *ℓ*(*t*) measured from the growth of single domains (Appendix A), we find that *f*(*x*) behaves similar to a power law over a wide range of *n*_*k*_, with an effective exponent between -0.65 and -0.85. Using the same rescaling as for the power-law growth for the measured *f*(*x*) gives reasonable collapse over a range of values of *u* and *L* (Fig. A6) with a power law decay *f*(*x*) ∼ *x*^−0.72^. We note that *f*(*x*) measured from simulations appears closer to a power-law form for *x* → 0 than the deterministic approximation.

**FIG. A6.**
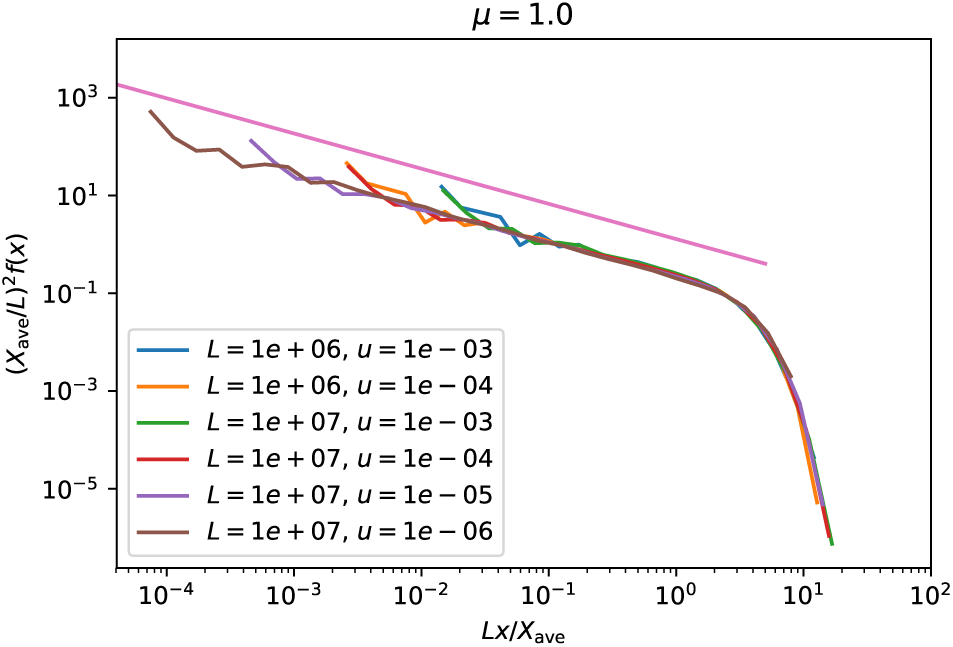
Comparison of allele frequency spectra over a range of system sizes for *µ* = 1. Allele frequency spectra for *µ* = 1 over different values of *L* and *u*, rescaled according to the assumption that the only length scale for the domains is *χ*. The low-frequency behaviour is consistent with a powerlaw decay that goes as *x*^−0.72^ (straight line).

### 3. Stretched exponential growth

In the stretched-exponential growth regime *µ* < *d*, the rescaling of the frequency spectra for a specific kernel proposed in Equation 4 is no longer exact. The rescaling assumed that *χ* set all length scales in the problem; this was true for power-law growth because the halo-dependent scales *ψ* and *ζ* were proportional to *χ* (with proportionality factors that depended only on *µ* and not on *χ*). By contrast, for stretched-exponential growth the additional length scales depend on the average clone sizes and hence on *ũ*. However, Fig. 6 showed that the rescaling captured much of the variation in *f*(*x*) across two well-separated mutation rates, down to *µ* = 0.4.

Although we could compute approximate frequency spectra using the deterministic calculation outlined above, they are less revealing in this regime. Instead, we gauge the inaccuracy of the proposed scaling in the panmictic limit *µ* → 0 where we know the exact frequency spectrum *f*_∞_. When *Nũ*= *θ* ≫ 1, we have *X*_ave_ ≈ – 1/(*ũ* log *ũ*) in the panmictic limit. Using this result and the form for *f*_∞_ in Eq. 4, we find that

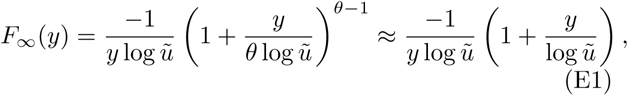

where *y* = *N*_*x*_/*X*_ave_ and we have used *θ* ≫ 1 in the second step. We find that the function after rescaling has a residual dependence on log *ũ*, both in the overall magnitude and in the value *y*_c_ ∼ log*ũ* of the dropoff in *f*. The gentle logarithmic correction implies that the proposed rescaling still captures much of the variation with mutation rates for a given kernel, even ifũis varied by orders of magnitude, thus explaining the decent collapse of curves at different mutation rates in Fig. 6 even for *µ* < *d*.

## Appendix F: Allele frequency spectra with a hard cutoff

The measured allele frequency spectra display a powerlaw behaviour *f*(*x*) ∼ *x*^*p*^, *p* > – 1 for small values of *x.* For cores growing as contiguous domains, balancing growth and mutation rates gives rise to a characteristic linear domain size *χ* (and corresponding clone size *χ*^*d*^) for domain growth before a cone encounters a new mutation. In a finite range of size *L*^*d*^, such growth would imply an upper bound on the allowed allele frequency at some value *x*_c_ ∼ (*χ*/*L*)^*d*^. These observations suggest the ansatz for the allele frequency spectra introduced in the main text:

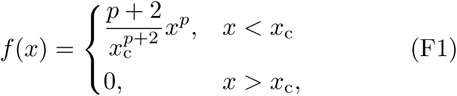

where the prefactor is determined by the normalization condition 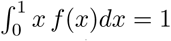.

This ansatz ignores contributions from higher-frequency clones, which are clearly significant especially for small values of *µ*. We can evaluate the significance of these contributions by comparing measured quantities to expectations from the hard-cutoff ansatz below.

The average clone size *X*_ave_ ≡ *N*/*n*_*c*_ = *N*/ ∫ *f*(*x*)*dx* can be evaluated for all *p* > −1 as

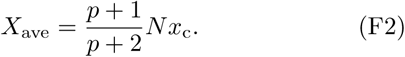

The sampling probability of observing only one allele in a sample of size *j* evaluates to

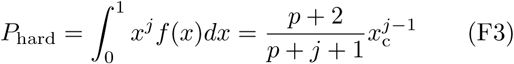

which deviates weakly from the exponential falloff *P*_hard_ = *x*^\**j*–1^ expected if all clones are of the same size and hence the same frequency *x*^*^.

## Appendix G: Sampling statistics in panmictic and 1D wavelike growth limits

In the panmictic limit, *µ* → 0, sampling probabilities are known analytically for all sample sizes [18]. Using *f*_*∞*_(*x*) in Eq. 6 gives *P*_hard_ = *θ*(*j* – 1)!Γ(*θ*)/Γ(*j* + *θ*) [5, 18] (where Γ denotes the gamma function). The result has two distinct behaviours depending on the value of *θ* = *N ũ*. When *θ* ≫ 1, an exponential falloff *P*_hard_ ∼ (1/*θ*)^*j*^ *θ*Γ(*θ*) is recovered for large *j*, whereas for *θ* ≪ 1, *P*_hard_(*j*) falls slower than 1 – *θ* log *j*.

For 1D wavelike growth with constant front velocity, Ref. [19] provides the exact form for the allele frequency spectrum, Eqs. D2–D3. The probability of observing only one allele in a random sample of size *j* is then 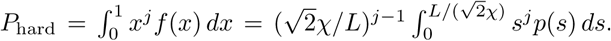. The latter integral cannot be evaluated in a closed form, even when we consider *L*/*χ* ≫ 1 so that the upper limit can be replaced by *s* = ∞ However, by tracking the position of the maximum value of the integrand which occurs at 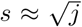, and using Laplace’s method to approximate the integral, we arrive at 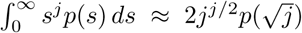, wh<u>i</u>ch provides a correction to the leading contribution 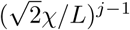. The resulting approximate expression,

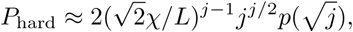

is used in the dash-dotted line in Fig 7 of the main text. Note that the approximation is only valid when the maximum value of the integrand lies below the upper integration limit; i.e. for *j* < *L*^2^/(2*χ*^2^). For larger values of *j*, *P*_hard_ is dominated by the upper limit, and scales as 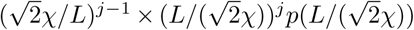 which is independent of *j*; i.e. the probability of detecting a hard sweep ultimately levels off for sufficiently large *j*.

